# Combinatorial characterization of bacterial taxa-driven differences in the microbiome of oyster reefs

**DOI:** 10.1101/2024.05.15.594453

**Authors:** Erika L. Cyphert, Sanjiev Nand, Gabriela Franco, Michael Hajkowski, Luzmaria Soto, Danica Marvi Lee, Matt Ferner, Chela Zabin, Jeffrey Blumenthal, Anna Deck, Katharyn Boyer, Kai Burrus, Christopher J Hernandez, Archana Anand

## Abstract

Oyster reefs are invaluable ecosystems that provide a wide array of critical ecosystem services, including water filtration, coastal protection, and habitat provision for various marine species. However, these essential habitats face escalating threats from climate change and anthropogenic stressors. To combat these challenges, numerous oyster restoration initiatives have been undertaken, representing a global effort to preserve and restore these vital ecosystems. A significant, yet poorly understood, component of oyster reefs is the microbial communities. These communities account for a substantial proportion of marine reefs and are pivotal in driving key biogeochemical processes. Particularly, the environmental microbiome plays a crucial role in supporting the health and resilience of oyster populations. In our study, we sought to shed light on the microbiome within oyster reef ecosystems by characterizing the abundance, and diversity of microorganisms in the soil, biofilm, and oysters in 4 sites using a combinatorial approach to identify differentially abundant microbes by sample type and by sampling location. Our investigation revealed distinct microbial taxa in oysters, sediment and biofilm. The maximum Shannon Index indicated a slightly increased diversity in Heron’s Head (5.47), followed by Brickyard park (5.35), Dunphy Park (5.17) and Point Pinole (4.85). This is likely to be driven by significantly higher oyster mortality observed at Point Pinole during routine monitoring and restoration efforts. Interestingly *Ruminococcus, Streptococcus, Staphylococcus, Prevotella, Porphyromonas, Parvimonas, Neisseria, Lactococcus, Haemophilus, Fusobacterium, Dorea, Clostridium, Campylobacter, Bacteroides*, and *Akkermansia* were positively associated with the biofilm. Yet we have limited understanding of their beneficial and/or detrimental implications to oyster growth and survival. By unraveling the intricate relationships in microbial composition across an oyster reef, our study contributes to advancing the knowledge needed to support effective oyster reef conservation and restoration efforts.

## Background and Introduction

Oyster reefs are invaluable ecosystems, offering a myriad of ecosystem services that are fundamental to both environmental health and human well-being. One of the primary functions of oyster reefs is water filtration. Adult oysters are capable of filtering up to 50 gallons of water per day, removing particulates, nitrogen, and other pollutants, thus improving water quality and clarity (NOAA, 2020; Zu Ermgassen et al., 2015; Luckenbach et al.,1999). This filtration process is essential for maintaining the health of marine waters (Yu & Gan, 2021), controlling algal blooms (Jiang et al 2019), and providing clearer water, which supports the growth of submerged aquatic vegetation (Kellogg et al., 2014; Newell, 2004). Furthermore, oyster reefs serve as an effective form of coastal defense. The physical structure of the reefs reduces wave energy and protects shorelines from erosion (Morris et al., 2019; Scyphers et al., 2011). They serve as natural barriers, absorbing wave force and mitigating the potential impact of storm surges, a service that is particularly valuable in the face of increasing climate change-related extreme weather events (Perricone et al 2023). In addition to water filtration and coastal protection, oyster reefs also provide critical habitats for a diverse array of marine species. As biogenic structures, they create three-dimensional living spaces that are utilized by various organisms for breeding, feeding, and shelter (Coen et al., 2007; Chan et al., 2022). The complex habitat created by oyster reefs supports high biodiversity and can include fish, crustaceans, and other invertebrates, thereby contributing to the productivity and function of estuarine ecosystems (Grabowski et al., 2012; Grabowski et al., 2022). These habitats not only enhance fishery resources but also contribute to the overall resilience of marine ecosystems. Preserving and restoring oyster reefs is, therefore, not only vital for the ecosystem services they provide but also for maintaining biodiversity and ensuring the sustainability of coastal resources (Smith et al., 2023; Beck et al., 2011).

However, these essential habitats are increasingly imperiled by a confluence of threats originating from both climate change and a range of human activities. Climate change impacts, such as ocean acidification and rising sea temperatures, have been shown to affect the growth and survival of oyster populations (Wilberg et al., 2011; Waldbusser et al., 2015). For instance, alterations in ocean chemistry due to increased carbon dioxide absorption can compromise the oysters’ ability to build and maintain their calcium carbonate shells, a process critical for their survival (Gledhill et al., 2015). Additionally, Okon et al (2023) in a global review highlight how aquatic pathogens promoted by climate change will likely affect oysters. Anthropogenic stressors provide an additional burden to oyster reefs. Coastal development often leads to habitat destruction, while pollution and overharvesting further exacerbate the decline of these ecosystems (Beck et al., 2011; Coen & Luckenbach, 2000). The deterioration of water quality due to nutrient run-off, for example, can lead to hypoxic conditions detrimental to oysters and the myriad of species that depend on oyster reefs (Dame & Dame, 1996).

In response to these escalating challenges, numerous oyster restoration initiatives have emerged globally, showcasing a concerted effort to preserve and restore these vital ecosystems (Beck et al., 2009). Such efforts often involve collaborations among scientists, resource managers, and stakeholders, employing strategies from reseeding programs to the installation of artificial reefs designed to mimic the function of natural oyster beds (Brumbaugh & Coen, 2009; Baggett et al., 2015; Perricone et al., 2023). These initiatives not only aim to bolster declining oyster populations but also to reestablish the ecosystem services that have been compromised (Ridlon et al., 2021; Schulte et al., 2014; Boudreau & Worm, 2012). Restoration projects such as those conducted in the Chesapeake Bay, USA, and the Solent, UK, have demonstrated success, with restored reefs providing critical ecological functions and economic benefits alike (Coen et al., 2007; zu Ermgassen et al., 2016). It is evident that these restoration projects are integral components of a larger strategy for marine conservation and are increasingly recognized for their role in supporting the resilience of coastal systems to future environmental changes (Grabowski et al., 2012).

A significant, yet poorly understood, component of oyster reefs is the intricate mosaic of microbial communities. Microbiomes form a substantial proportion of biofilm on marine reefs and are pivotal in cycling nutrients and transforming materials through key biogeochemical processes (Dame, 1996; Dang & Lovell, 2016; Remple et al., 2021; Mannochio-Russo et al., 2023). The microbial elements within oyster reefs undertake roles from nitrogen fixation to sulfate reduction, which are critical in the maintenance of water quality and overall ecosystem health (Welsh et al., 2016). Particularly, the environmental microbiome plays a crucial role in supporting the health and resilience of oyster populations. These microbial assemblages are involved in the breakdown of organic matter, which ensures the availability of nutrients for oyster growth, and contributes to the stabilization of the reef structure (Freeman et al., 2016). Moreover, oysters interact with their microbiomes for key physiological processes, including digestion, immunity, and possibly even adaptation to stressors like those presented by climate change (King et al., 2012). Symbiotic relationships between oysters and certain bacteria can enhance the bivalves’ defense mechanisms against pathogens, reducing disease incidence and improving survival rates, thus implying an intricate connection between microbial communities and oyster resilience (Acevedo-Whitehouse & Duffus, 2009; Trabal et al., 2012; Newton et al., 2007; Sakowski et al., 2020). Intriguingly, research has revealed that alterations in the microbial community composition can be indicative of changes in oyster health and environmental quality, signifying the potential for these communities to serve as biological indicators (Trabal Fernandez et al., 2014).

Despite their importance, the complexity of microbial communities and their interactions with environmental factors and host species are not fully understood. Recognizing the extent to which these dynamic communities contribute to oyster reef ecosystems is essential for their management and conservation (Lokmer & Wegner, 2015). Enhanced understanding can inform restoration efforts, ensuring that the environmental conditions support both the oysters and their associated microorganisms, which collectively facilitate the functioning and persistence of these vital habitats (Lokmer & Wegner, 2015; Baggett et al., 2015).

In our study, we sought to shed light on the microbiome within oyster reef ecosystems by characterizing the abundance and diversity of microorganisms present in the soil, biofilm, and oysters collected from reefs with varying oyster population densities in San Francisco Bay. Understanding the dynamics of microbial populations in these habitats is crucial, as microbial diversity and abundance can be indicators of ecosystem health and function (Allison & Martiny, 2008). We utilized 16S rRNA gene sequencing, to provide a comprehensive profile of the microbial communities across different components of the oyster reef ecosystem (Caporaso et al., 2012; Burgess et al., 2017). By quantifying and comparing microbial communities across a gradient of oyster densities, our analysis aimed to answer the research question - how does the microbiome of oysters, biofilm and soil vary?

## Materials and Methods

### Site Description

The sampling strategy included a systematic collection from four sites (Point Pinole, Brickyard Park, Dunphy Park, Heron’s Head) with varying oyster densities in San Francisco Bay in November 2023 (**Fig.1**), which enabled an examination of the potential correlations between oyster populations and microbial diversity. Soil samples were analyzed to understand the baseline microbial communities within the reef structure, while biofilm samples were used to assess the microorganisms directly associated with the oyster’s surface—critical for oyster health and nutrient cycles within the reef (Harris et al., 2016). Oyster tissue samples were also collected and analyzed to investigate the gut microbiome, which plays a fundamental role in the oyster’s digestion and immune function (King et al., 2012). By considering these distinct but interconnected microbial habitats, we intended to capture a holistic view of the microbiome’s role within the oyster reef ecosystem and assess the implications for oyster reef conservation and restoration efforts.

**Figure 1.**
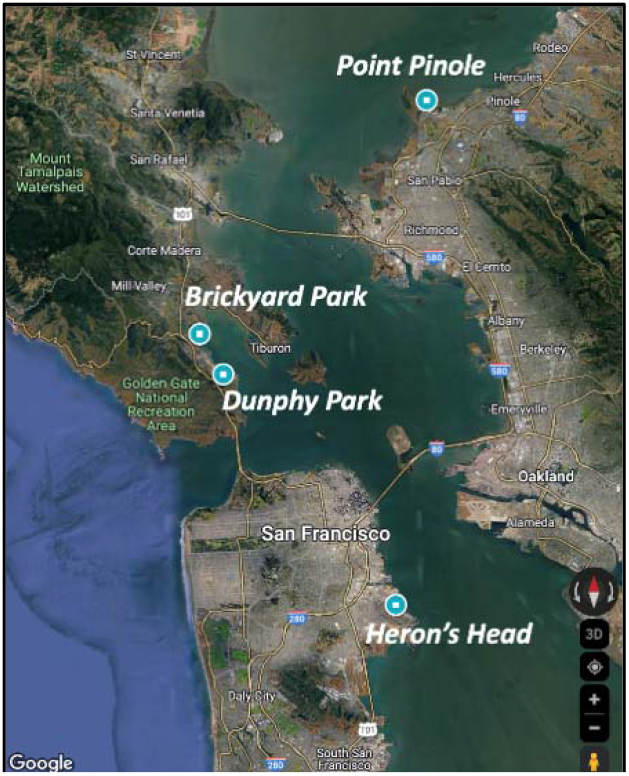
Study sites for oyster, biofilm and soil sampling in San Francisco Bay

### Sample Collection

To collect sediment, three 50 mL falcon tube were filled with sediment from the top 10 cm from every oyster sampling site, following which they were placed in a sterile bag. To collect biofilm, three 15mL falcon tubes were first filled with 8-10 mL of 99.5% ethanol. Using a sterilized scraper, biofilm from substrates were scraped and placed into the ethanol-filled tube while taking care to not disturb barnacles or other sessile organisms present. To collect oysters, five or more oysters were collected per site and placed in separate sterilized bags (WhirlPak). All collected samples were stored in a cooler for transportation to San Francisco State University campus, where they were subsequently stored in a -80°C freezer for downstream processing.

### Concentration and Nucleic Acid Extractions

For oyster tissue DNA extraction the DNeasy Blood & Tissue Kit (Qiagen) was used following the manufacturer’s instructions. 25 mg ± 1 mg of oyster tissue was used. The samples were incubated in a water bath at 56°C overnight (12+ hrs.). For maximum yield, the elution step was repeated as per the manufacturer’s protocol. For soil, and biofilm DNA extractions the DNeasy PowerSoil Pro Kit was used following the manufacturer’s instructions. Given that the biofilm was suspended in isopropyl alcohol (99.5%) upon retrieval, instead of a 250 mg sample being used 250 μl of the mixture was used. DNA extracts were stored in a -80°C freezer until they were shipped. DNA extracts were then sent for 16S rRNA amplicon sequencing to the Genomics Core at University of California San Diego. Before sequencing, the 16S rRNA V4-V5 variable region was amplified with primers 515F and 806R (Caporaso et al 2011) using Illumina MiSeq.

### Microbiome Analysis

Paired-end sequences (2 × 150 bp) were processed and taxonomically classified using QIIME2 (v. 2020.6) and SILVA database (SSU r138-1) (Estaki et al 2020; Cyphert et al 2023). Reads were normalized using a single rarefaction step (feature count cut-off: 6993) (Weiss et al 2017; Mallick et al 2017). Amplicon sequence variants (ASVs) generated in QIIME2 were used calculate alpha diversity (Shannon index; Richness) and beta diversity (Bray-Curtis dissimilarity) using the vegan package (v. 2.5-7) in R (v. 4.0.1) (Dixon 2003). A principal coordinate analysis (PCoA) was carried out on Bray-Curtis beta diversity (post-rarefy) and samples were grouped relative to sample type and sampling location (95% confidence ellipses). Differentially abundant genera by sample type (positively or negatively associated relative to Point Pinole) and sampling site (positively or negatively associated relative to Sediment) were determined using Microbiome Multivariate Associations with Linear Models (MaAsLin2) (Mallick et al 2021). MaAsLin output is represented in a heatmap where the intensity of association reflects -log_10_ transformed post-hoc q-values. Additionally, a secondary analysis using ALDEx2 R package was carried out on ASVs generating Monte Carlo simulations of Dirichlet distributions (centered log ratio transformed) for samples and a generalized linear model was determined based on sample type (Oyster versus Biofilm; Sediment versus Biofilm) (Fernandes et al 2014; Nearing et al 2022). ALDEx2 results are represented in volcano plots where significant ASVs are indicated in upper left and right corners (p < 0.05; magnitude of fold change > |1|).

### Statistical Analysis

Statistical analyses were carried out using RStudio (v. 1.4.1106, Boston, MA, US, 2021) (RStudio 2021). For alpha diversity metrics (Shannon index, Richness) a one-way analysis of variance (ANOVA) calculated significance across sample type and sample location. For Bray-Curtis beta diversity, a permutational multivariate analysis of variance (PERMANOVA; adonis2 function) calculated significance in microbiota composition by sample type and sample location (Anderson 2017).

### Data availability

Raw V4-V5 16S rRNA DNA sequences are available at the NCBI’s Sequence Read Archive Database (BioProject ID: PRJNA1108902; http://www.ncbi.nlm.nih.gov/bioproject/1108902)

## Results

Results showed 4919 unique OTUs in all samples. Bacterial OTUs were assigned to 55 different phyla while Archaea OTUs were assigned to 10 different phyla (*Crenarchaeota, Nanoarchaeota, Thermoplasmatota, Asgardarchaeota, Aenigmarchaeota, Altiarchaeota, Halobacterota, Hydrothermarchaeota, Hadarchaeota, Iainarchaeota*). Across all four sites, the dominant taxa were bacteria phyla *Proteobacteria, Bacteroidota, Desulfobacteria, Firmicutes* and *Cyanobacteria*. These findings are consistent with findings from literature that have utilized 16S to quantify relative abundances of individual taxa in different oyster sites (Steves et al 2019). The top 20 genera in biofilm revealed the presence of animal pathogens (unclassified genus within *Enterobacteriaceae, Streptococcus, Neisseria, Staphylococcus*) and naturally derived microorganisms with probiotic potential (e.g. *Lactococcus, Akkermansia, Alistipes*) that resist the colonization of undesirable genera. *Sulfuvorum* and *Fusobacterium* were noted to be present in oysters that were undergoing mortality in the natural environment and getting spoiled in the refrigerator respectively. The most dominant genus belonged to an Archaeal family - *Thermoplasmatota* (*Candidatus Nitrosopumilus*).

When divided by sample type, the Bray-Curtis beta diversity showed significant differences between Biofilm, Oyster, and Sediment samples (**Figure 2A**; PERMANOVA p = 0.002) with the Oyster and Sediment having more similar compositions than in the Biofilm. When divided by sampling location, there were significant differences in the microbiome composition (**Figure 2B**; PERMANOVA p = 0.022) with the biggest differences in the Point Pinole location. Shannon index and richness were the greatest in the Sediment relative to the Biofilm (Shannon: p = 0.017; Richness: p = 0.016) and Oyster (Shannon: p = 0.368; Richness: p = 0.062) (**Figure 2C,E**). The maximum Shannon Index indicated a slightly increased diversity in Heron’s Head (5.47), followed by Brickyard park (5.35), Dunphy Park (5.17) and Point Pinole (4.85) (**Figure 2D**). Richness also indicated a similar trend (485, 440, 496 and 386 at Heron’s Head, Brickyard Park, Dunphy Park and Point Pinole respectively (**Figure 2F**). We hypothesize that this could be attributed to significantly higher oyster mortality observed at Point Pinole during routine monitoring and restoration efforts, during our sampling period when compared to the other sites (unpublished data).

**Figure 2.**
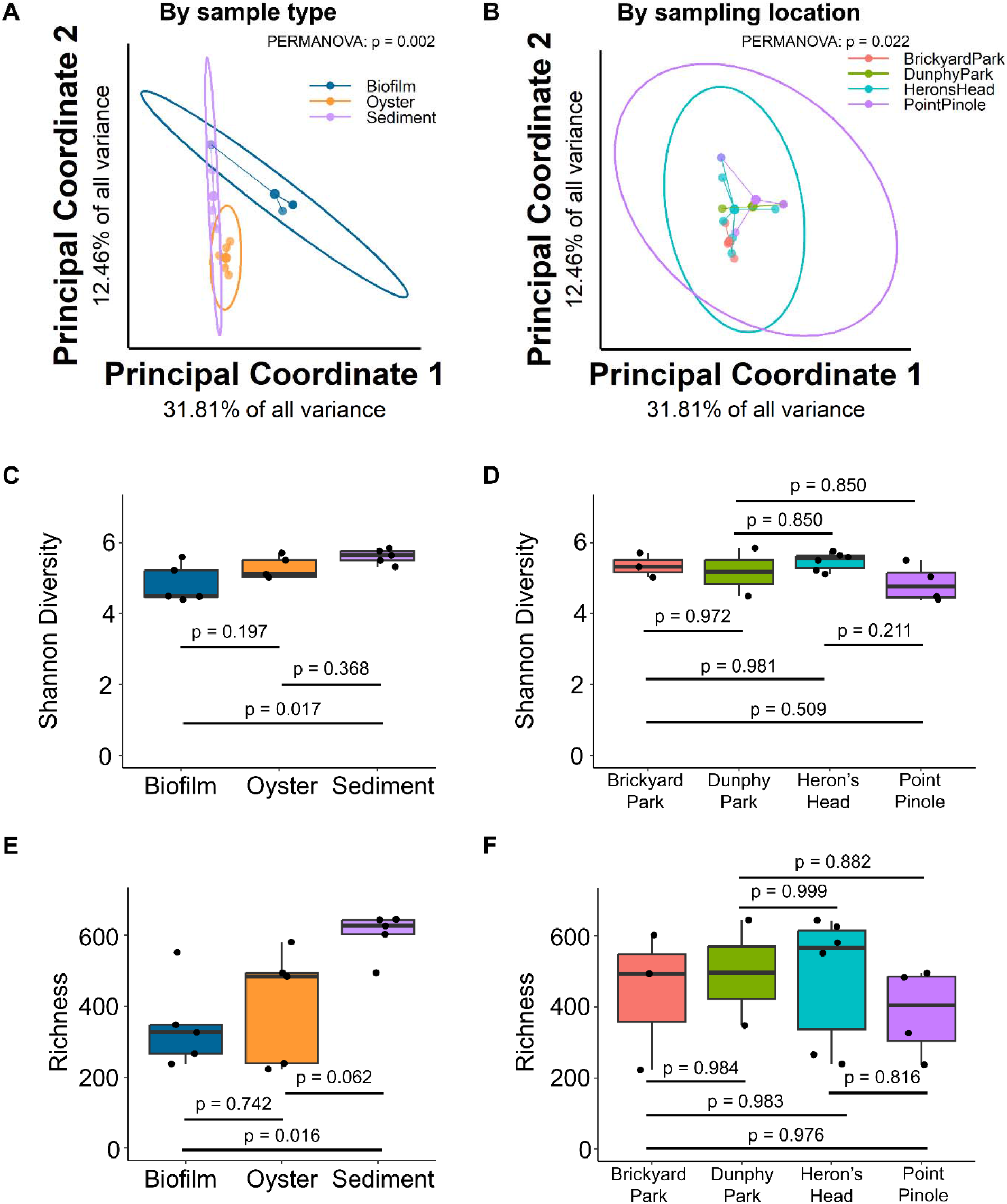
Differences in microbiota composition based on sample type (**A**) and sample location (**B**) using principal coordinate analysis Bray-Curtis beta diversity. Differences in alpha diversity (Shannon Diversity and Richness) based on sample type (**C, E**) and sample location (**D, F**).

**Figure 3.**
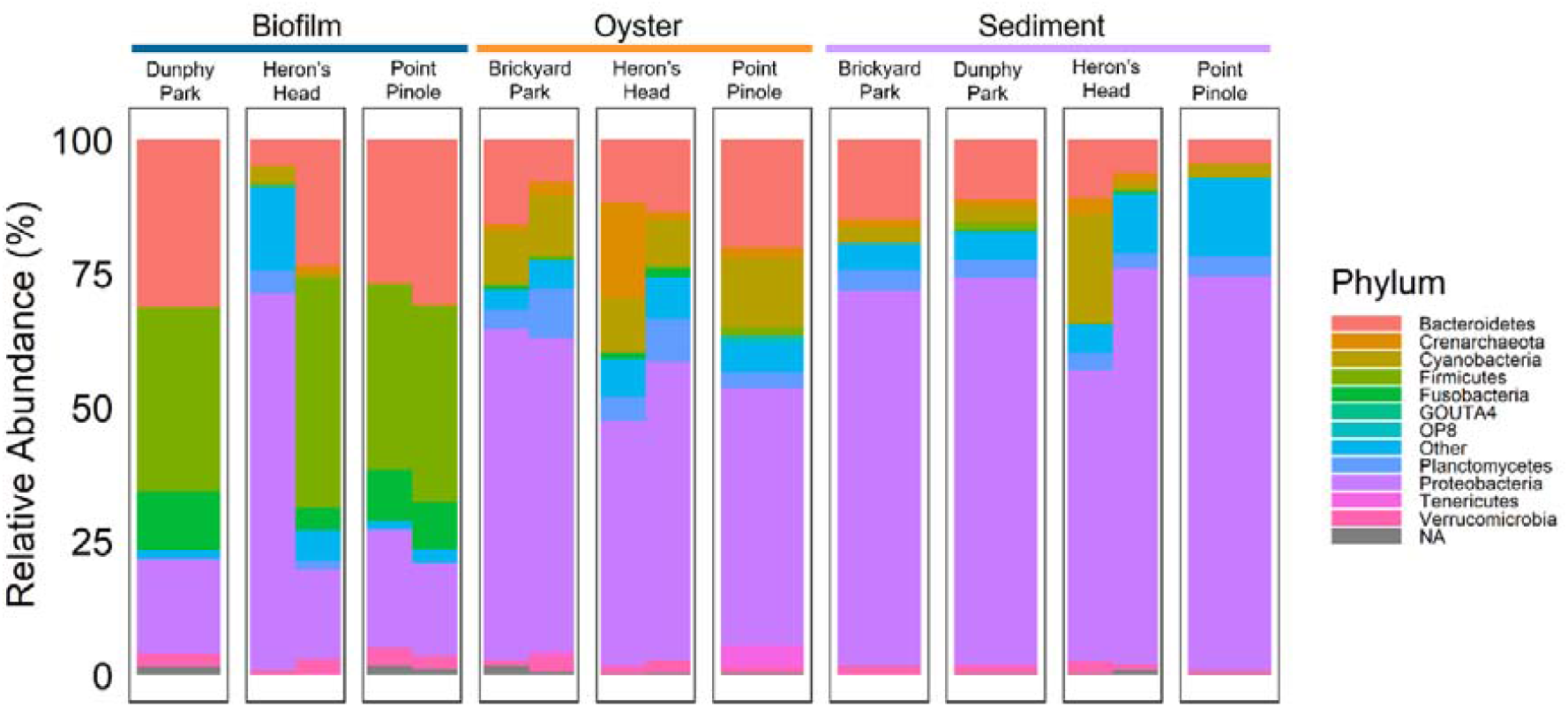
Relative abundance at phyla level based on sample type (Biofilm, Oyster, Sediment) and location (Dunphy Park, Heron’s Head, Point Pinole, Brickyard Park). Biofilm had elevated levels of *Firmicutes* across sampling locations and Oyster and Sediment had elevated levels of *Proteobacteria* across sampling locations.

Biofilm samples across sampling locations (Dunphy Park, Heron’s Head, Point Pinole) revealed intriguing ecological dynamics. Specifically, biofilm had increased *Firmicutes* and decreased *Proteobacteria* relative to oyster and sediment samples. A combinatorial approach was used to identify differentially abundant microbes by sample type and by sampling location. MaAsLin2 analysis (**Figure 4A**) identified 17 differentially abundant genera in the Biofilm relative to Sediment samples in which *Sulfitobacter* and *HTCC* were negatively associated with Biofilm and *[Ruminococcus], Streptococcus, Staphylococcus, Prevotella, Porphyromonas, Parvimonas, Neisseria, Lactococcus, Haemophilus, Fusobacterium, Dorea, Clostridium, Campylobacter, Bacteroides*, and *Akkermansia* were positively associated with Biofilm. Oyster was very similar to Sediment composition with only *Sulfitobacter* being negatively associated with the Oyster. Brickyard Park and Dunphy Park had one differential genera relative to Point Pinole with *planctomycete* and *Paludibacter* positively associated respectively. Results from ALDEx2 differential abundance analysis (**Figure 4B**) were confirmatory with 47 significant differentially abundant ASVs (41 ASVs with positive fold change, 6 ASVs with negative fold change; see taxonomic identity of ASVs – **Supplementary Table 1**). In the Biofilm relative to the Sediment there were 80 significant differentially abundant ASVs (45 ASVs with positive fold change, 35 ASVs with negative fold change; see taxonomic identity of top 50 ASVs – **Supplementary Table 2**). Between the Oyster and the Sediment there were 16 significant differentially abundant ASVs (1 with positive fold change and 15 with negative fold changes – **Supplementary Table 3**). Across the sampling sites significant differentially abundant ASVs were only detected between Brickyard Park relative to Point Pinole (8 ASVs with positive fold change – **Supplementary Table 4**), Brickyard Park relative to Heron’s Head (5 ASVs with positive fold change – **Supplementary Table 5**), and Brickyard Park relative to Dunphy Park (1 ASV with positive fold change – **Supplementary Table 6**).

**Figure 4.**
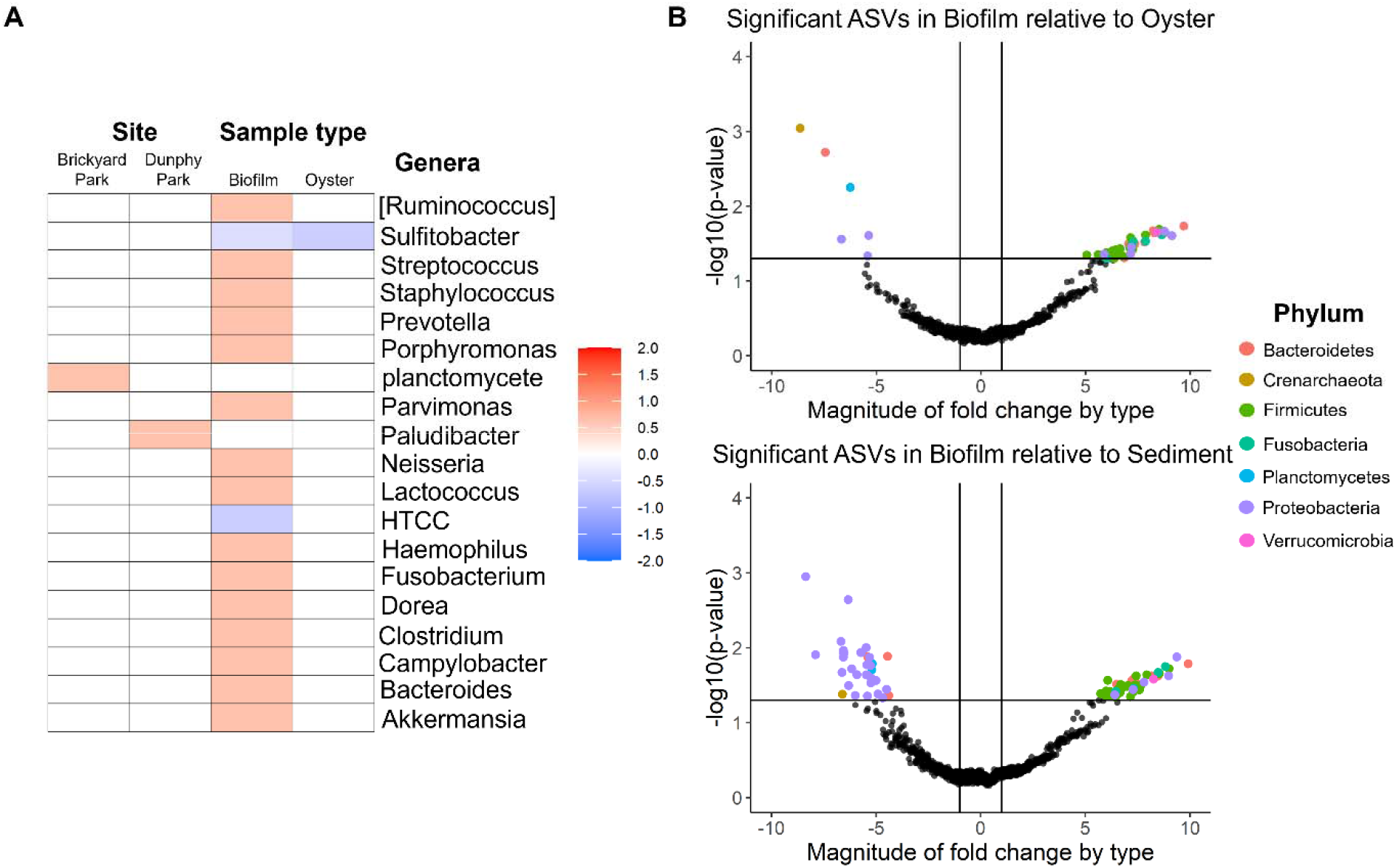
Differential abundance of microbes based on sample type and sampling site using two independent methods (MaAsLin2 – **A**; ALDEx2 - **B**). Genera positive and negatively associated with sampling site (Brickyard Park or Dunphy Park) relative to Point Pinole and associated with sample type (Biofilm or Oyster) relative to Sediment ASVs significantly increased or decreased in Biofilm relative to Oyster (top - **B**) or Sediment (bottom - **B**). Taxonomic classification of significant ASVs at the phyla-level is indicated.

## Discussion

### Functional specialization

The presence of increased *Firmicutes* and decreased *Proteobacteria* in biofilms within the sampled oyster reefs can be attributed to several factors, each potentially linked to the unique environmental conditions and biological interactions in these ecosystems (Trabal et al 2014). For example, biofilms in oyster reefs often experience low oxygen conditions due to the dense packing of organisms and the high biological activity. Firmicutes are known to thrive in anaerobic or low-oxygen environments, contributing to processes like fermentation and sulfate reduction, which are crucial for nutrient cycling in such ecosystems. *Firmicutes*, particularly some *Clostridia* species, are efficient at decomposing organic material in anaerobic or microaerophilic conditions, converting it into simpler compounds that can be utilized by other microorganisms or directly by oysters. Additionally, *Firmicutes*, particularly those forming spores, can withstand harsh environmental conditions, such as changes in salinity, temperature, and pH. Their presence in biofilms can enhance the resilience of the microbial community against environmental fluctuations. We hypothesize that challenging environmental conditions characterized by fluctuating salinity owing to severe atmospheric river events would’ve triggered the abundance of spore forming species. On the other hand, other microbial groups in the biofilm might outcompete *Proteobacteria* for nutrients or space. Additionally, bacteriophages or protozoan predators in biofilms might preferentially target *Proteobacteria*, reducing their numbers. Lastly, fermentation processes carried out by *Firmicutes* can produce organic acids and alcohols, creating a more acidic or toxic environment for some Proteobacteria.

### Physical structure and habitat

The significant differences in Bray-Curtis beta diversity among Biofilm, Oyster, and Sediment samples highlight distinct microbial community structures associated with each sample type. This suggests that the physical and biochemical environment of each sample type selectively enriches for different microbial assemblages. Notably, the greater similarity between Oyster and Sediment samples compared to Biofilm might indicate shared environmental conditions or an exchange of microbes between these habitats, possibly due to their proximity or similar physical characteristics. Additionally, differential abundance of certain microbes between locations such as Brickyard Park and Point Pinole, with genera like *Planctomycete* and *Paludibacter* positively associated respectively, suggests microbial adaptation to local environmental conditions or responses to anthropogenic effects (Diner et al 2023). Notably, Guedes et al (2023) demonstrated the ecological role of Planctomyceta and their antimicrobial properties.

### Detrimental vs. beneficial organisms

Interestingly, we observed the presence of a probiotic microbe in oysters (*Lactobacillales;* Kang et al 2018) in the biofilm relative to oyster and sediment. This suggests its role as a biocontrol agent against pathogenic microorganisms (Gatesoupe 2008). Notably, a recent study by Stevvick et al (2019) demonstrated that probiotic treatment in an oyster hatchery had a systemic effect on targeted members of the bacterial community, leading to a net decrease in potentially pathogenic species. The study also identified differentially abundant microbes, with *Sulfitobacter* and other genera showing distinct associations with Biofilm and Sediment samples. The negative association of *Sulfitobacter* with Biofilm and Oyster samples suggests its adaptability or preference for conditions present in Sediment. Bruhn et al (2007) demonstrated the antimicrobial capacity of strains such as *Sulfitobacter* belonging to the *Roseobacter* clade that produce antibacterial compounds that inhibit non-*Roseobacter* clade microbes and enhance biofilm formation.

On the other hand, Brickyard park had an increased abundance of both *Vibrio* and *Roseovarius* (both known to be pathogenic for oysters; Green et al 2019, Boardman et al 2008) relative to Point Pinole and Heron’s Head. Furthermore, the presence of specific genera in the Biofilm, such as *Streptococcus, Staphylococcus*, and *Akkermansia*, highlights unique microbial signatures that may be functionally detrimental or important in these environments. For example, Akkermansia’s current range is restricted to aquatic environments such as the human gut and is well known for its role in breaking down mucin and other glycoproteins. Our hypothesis is that Akkermansia plays an important role in regulating the presence of mucus that accumulates on the oyster substrates. This could have implications for understanding biofilm-associated diseases or health, presenting a targeted area for further ecological or biomedical research.

### Ammonia-oxidizing archaea

Ammonia-oxidizing archaea (AOA) are among the most ubiquitous microorganisms in the ocean, driving nitrification, nitrogen oxides emission and methane production.

Recent studies (Qin et al 2020) have demonstrated their adaptive capacity for nutrient acquisition and energy conservation through genetic diversification associated with niche adaptation. In our study sites, we observed significantly lesser AOA in biofilm than other sample types. We hypothesize that this is owing to their predominant role in nitrogen cycling by regulating the fixed forms of nitrogen species available (Li et al 2018; Martens-Habbena et al 2015) to other microbes present in the biofilm matrix and their adaptability to the available form of nitrogen present. AOA are more resistant to low-oxygen environments such as those in the gut of oysters and sediment, where they were also relatively more abundant than in biofilm. This suggests a vertical segregation of the AOA communities, consistent with literature (Lu et al 2016).

These findings could be further explored for their potential in ecological monitoring and assessment, contributing to our understanding of microbial roles in ecological processes like nutrient cycling and biofilm formation. These results collectively illustrate the complex interplay between microbial communities and their environments, highlighting the potential of microbial analyses in environmental monitoring and management strategies.

## Funding, Acknowledgments

We gratefully acknowledge the support of College of Science & Engineering, Department of Biology at San Francisco State University, National Science Foundation (NSF) Center for Cellular Construction, NSF Grant No. DBI-1548297, NIH R01AG067997 (C.J.H.), NIH F32AG076244 (E.L.C.), and the Chan Zuckerberg Biohub.

---

**Supplementary Table 1.**
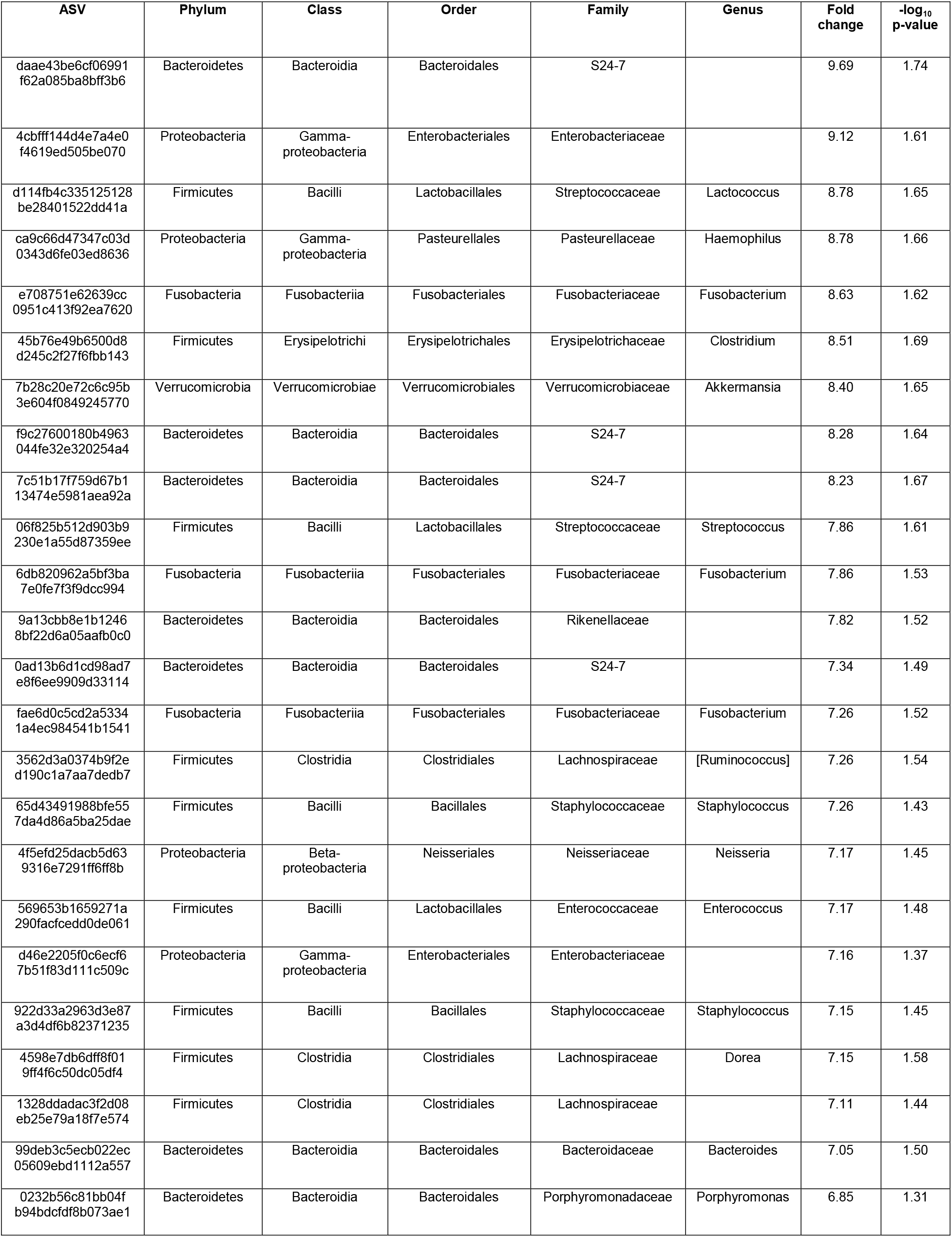
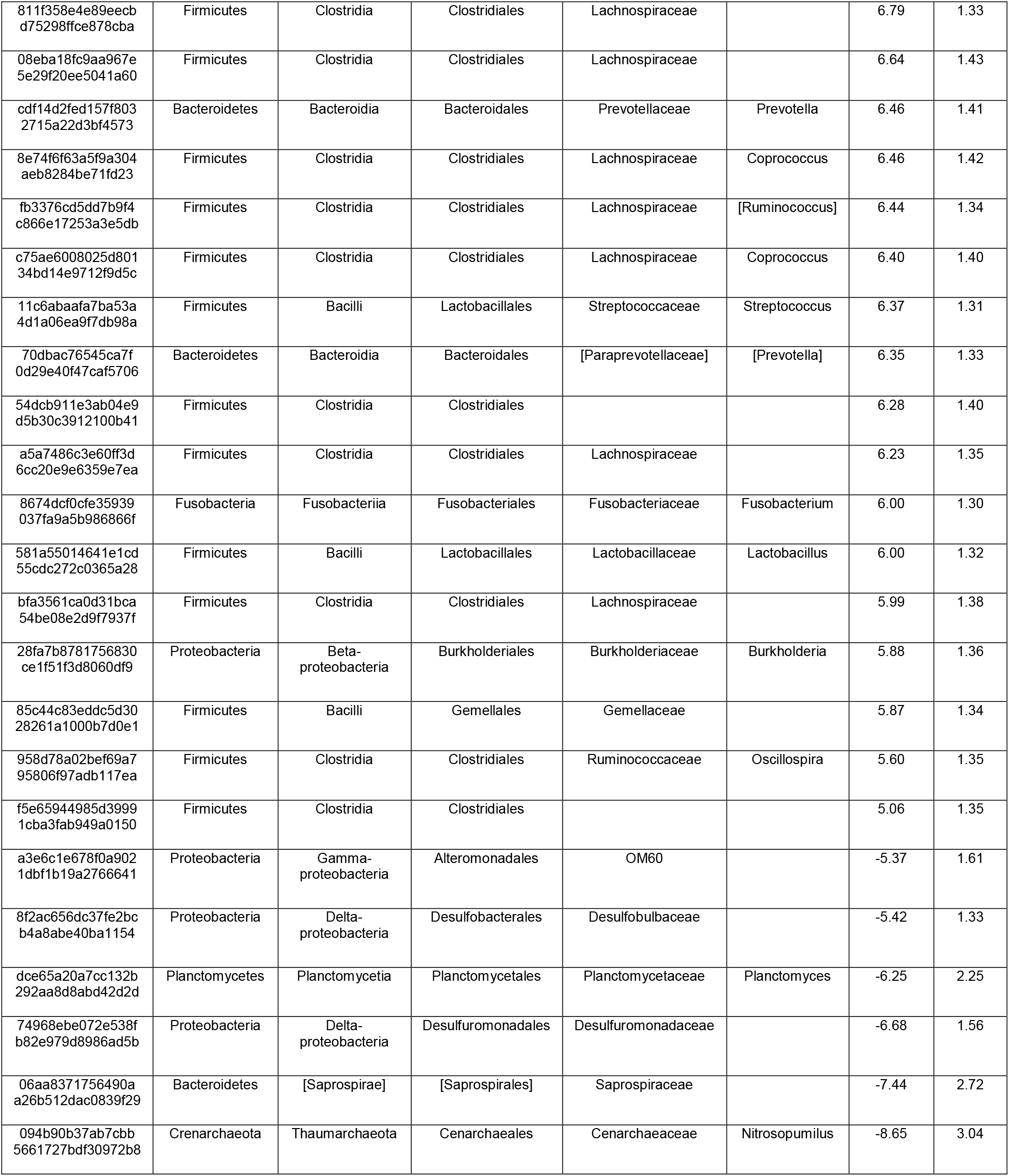
Taxonomy of significant ASVs (p < 0.05; magnitude of fold change > |5|) in Biofilm relative to Oyster identified using ALDEx2.

**Supplementary Table 2.**
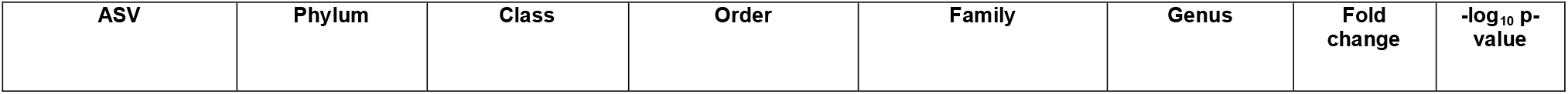
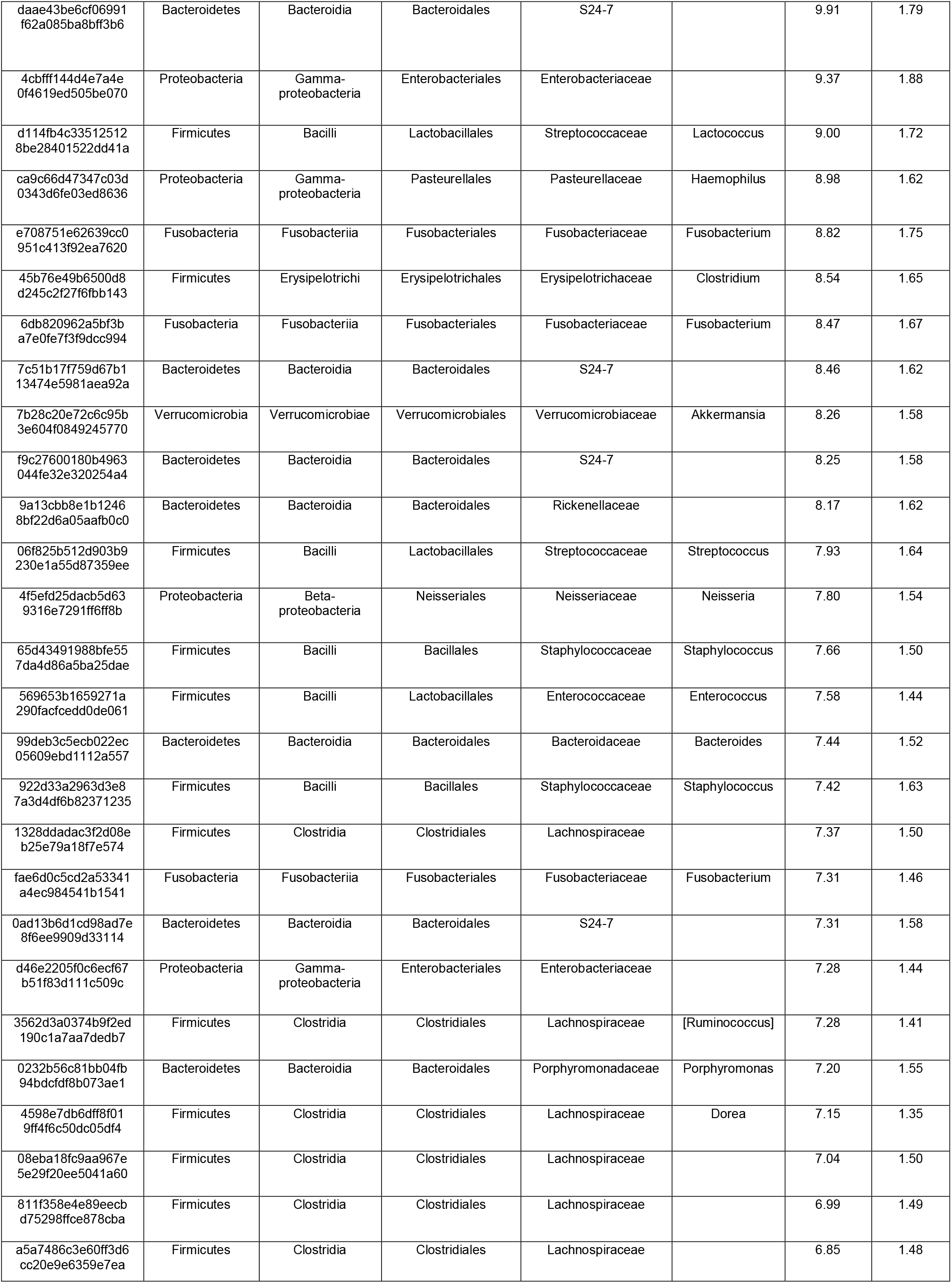
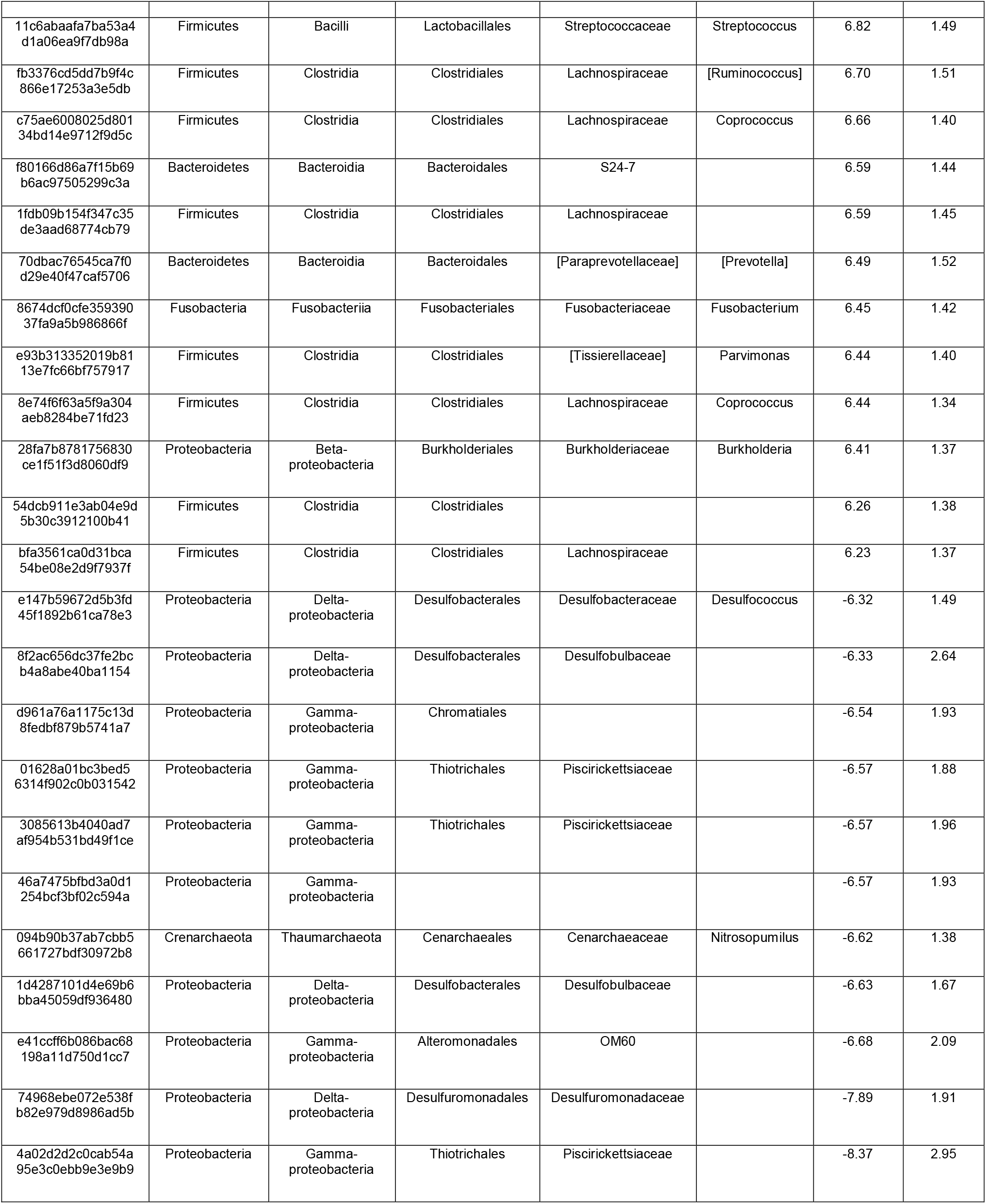
Taxonomy of top 50 significant ASVs (p < 0.05; magnitude of fold change > |6|) in Biofilm relative to Sediment identified using ALDEx2.

**Supplementary Table 3.**
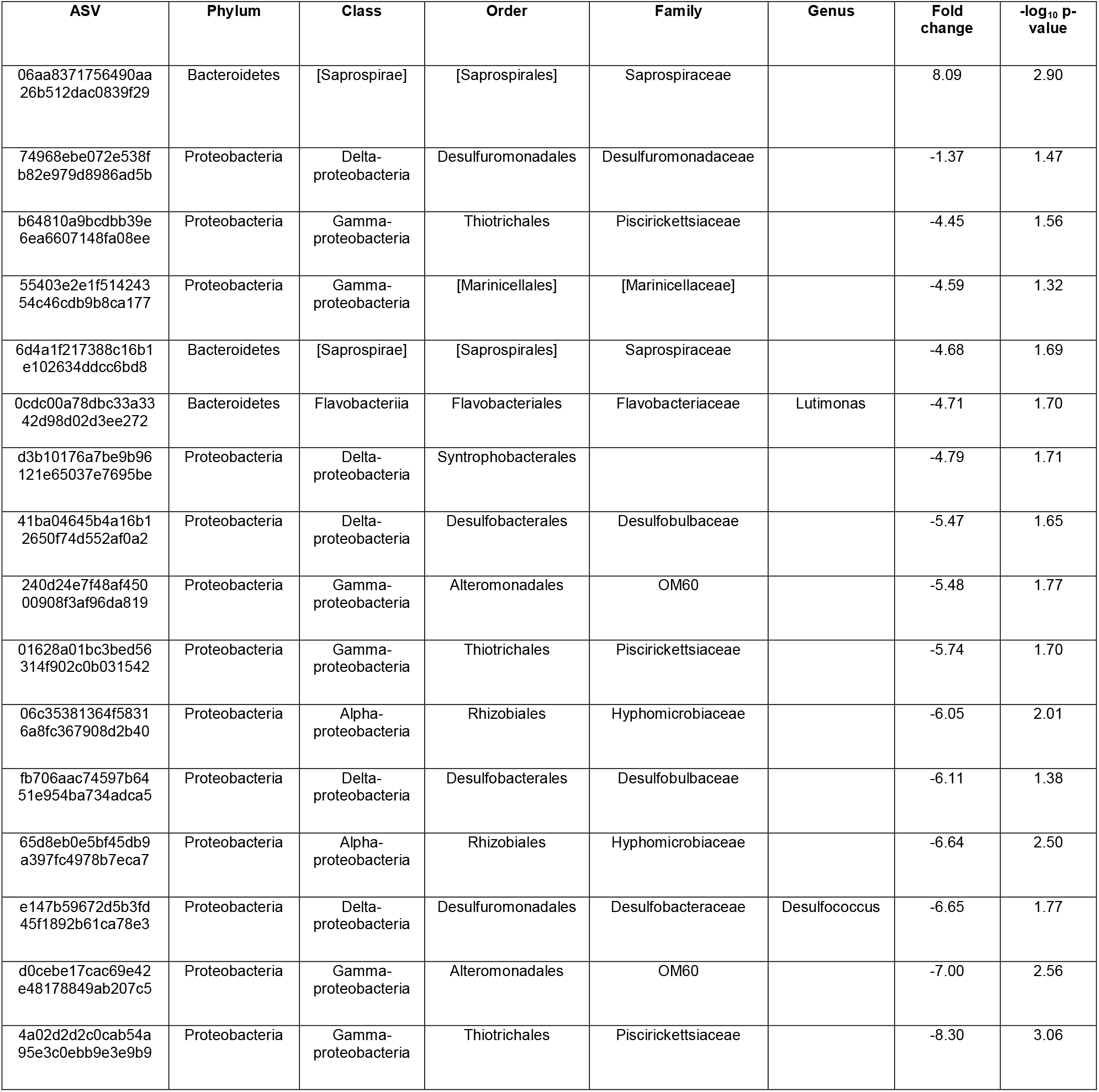
Taxonomy significant ASVs (p < 0.05; magnitude of fold change > |1|) in Oyster relative to Sediment identified using ALDEx2.

**Supplementary Table 4.**
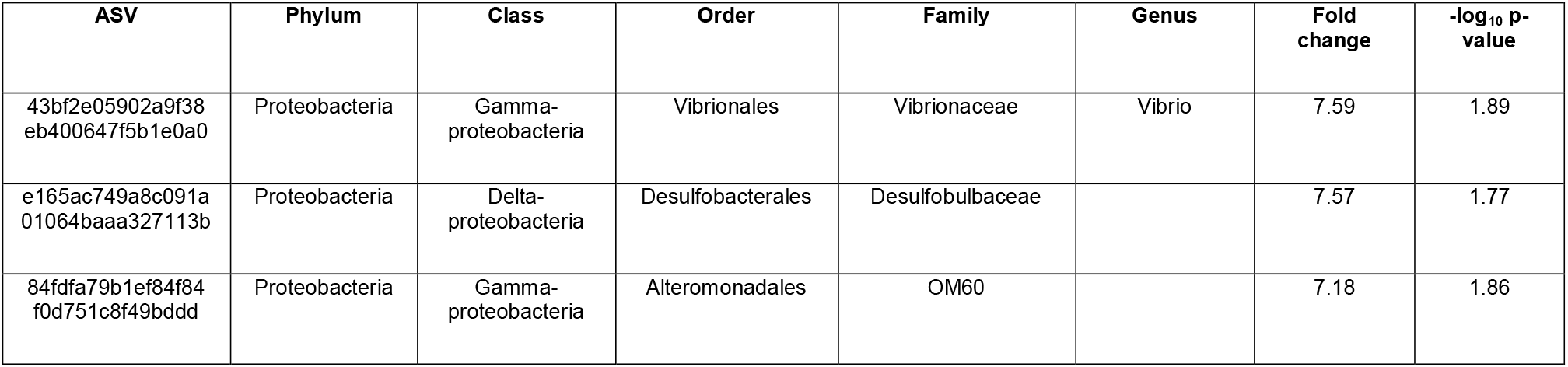
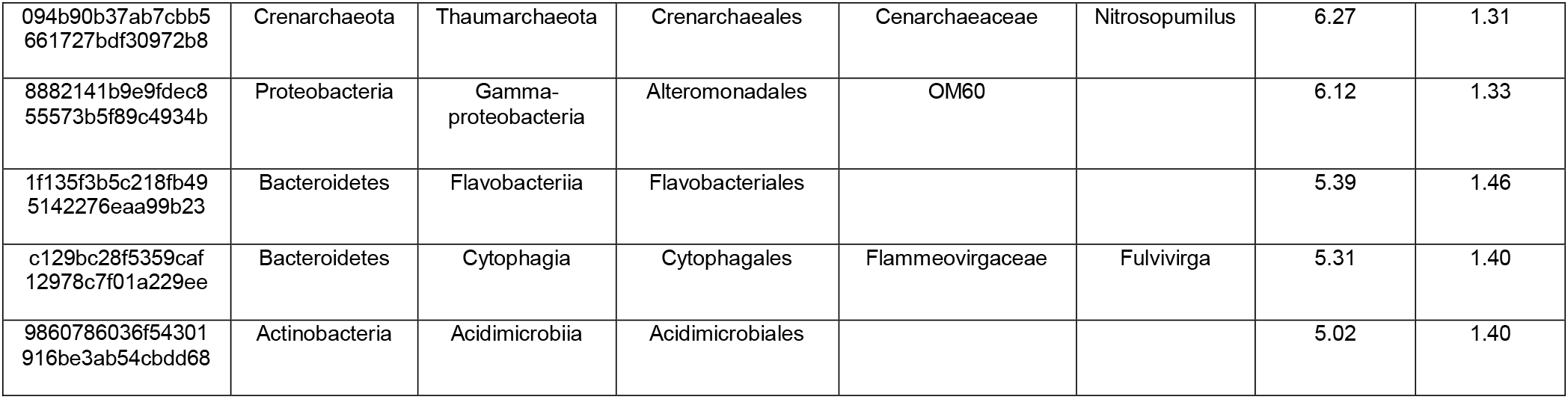
Taxonomy significant ASVs (p < 0.05; magnitude of fold change > |1|) in Brickyard Park relative to Point Pinole identified using ALDEx2.

**Supplementary Table 4.**
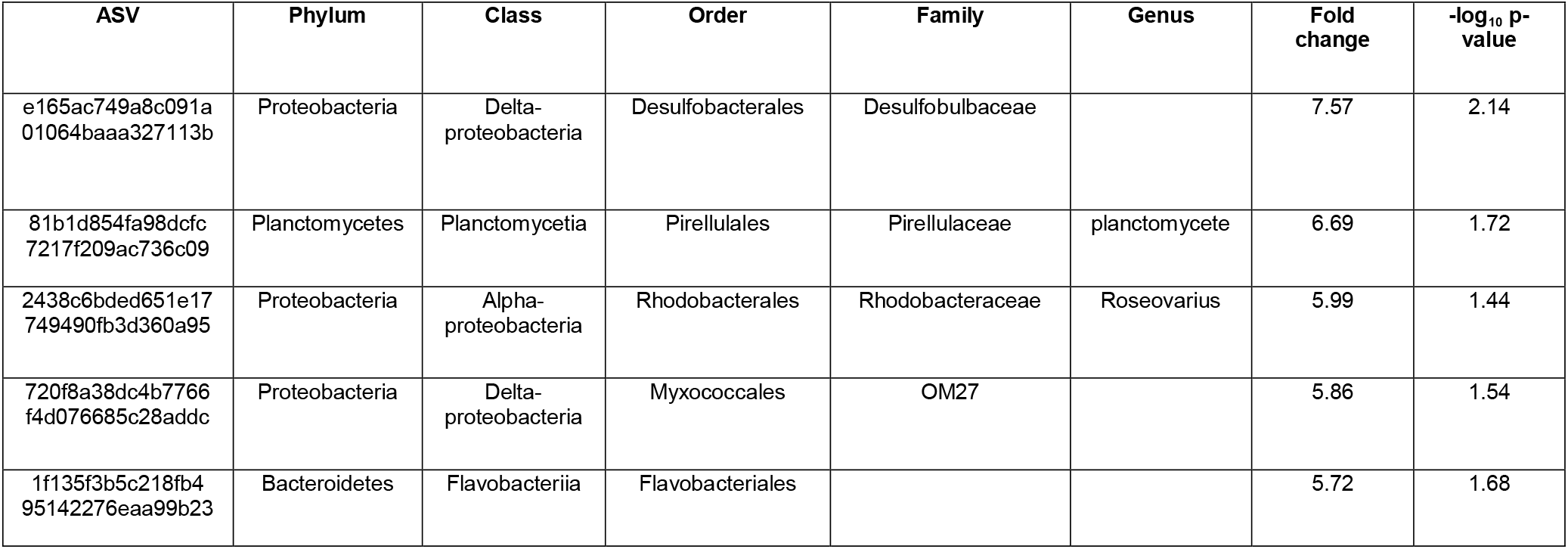
Taxonomy significant ASVs (p < 0.05; magnitude of fold change > |1|) in Brickyard Park relative to Heron’s Head identified using ALDEx2.

**Supplementary Table 5.**
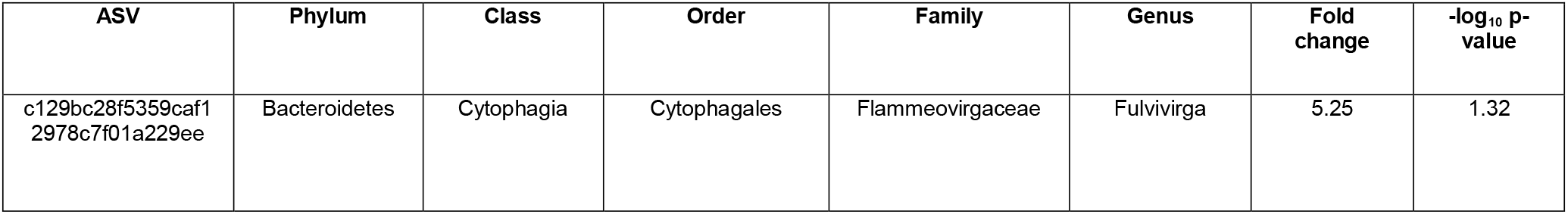
Taxonomy significant ASVs (p < 0.05; magnitude of fold change > |1|) in Brickyard Park relative to Dunphy Park identified using ALDEx2.

## References

1. Acevedo-Whitehouse, K., & Duffus, A. L. J. (2009). Effects of environmental change on wildlife health. Philosophical Transactions of the Royal Society B: Biological Sciences, 364(1534), 3429–3438.

2. Allison, S. D., & Martiny, J. B. H. (2008). Resistance, resilience, and redundancy in microbial communities. Proceedings of the National Academy of Sciences, 105(Supplement_1), 11512–11519.

3. Anderson M. Permutational Multivariate Analysis of Variance (PERMANOVA). Wiley StatsRef Stat Ref Online. 2017.

4. Baggett, L. P., Powers, S. P., Brumbaugh, R. D., Coen, L. D., Deangelis, B. M., Greene, J. K., Hancock, B. T., Morlock, S. M., Allen, B. L., Breitburg, D. L., Bushek, D., Grabowski, J. H., Grizzle, R. E., Grosholz, E. D., La Peyre, M. K., Luckenbach, M. W., Mcgraw, K. A., Piehler, M. F., Westby, S. R., & zu Ermgassen, P. S. E. (2015). Guidelines for evaluating performance of oyster habitat restoration. Restoration Ecology, 23(6), 737–745. 10.1111/rec.12262

5. Beck, M. W., Brumbaugh, R. D., Airoldi, L., Carranza, A., Coen, L. D., Crawford, C., Defeo, O., Edgar, G.J., Hancock, B., Kay, M.C., Lenihan, H.S., Luckenbach, M.W., Toropova, C.J., Zhang, G., Guo, X. (2011). Oyster reefs at risk and recommendations for conservation, restoration, and management. BioScience, 61(2), 107–116.

6. Beck, M.W., Brumbaugh, R.D., Airoldi, L., Carranza, A., Coen, L.D., Crawford, C., Defeo, O., Edgar, G.J., Hancock, B., Kay, M. and Lenihan, H. (2009) Shellfish reefs at risk. A global analysis of problems and solutions. Arlington, VA: The Nature Conservancy.

7. Boardman CL, Maloy AP, Boettcher KJ. Localization of the bacterial agent of juvenile oyster disease (Roseovarius crassostreae) within affected eastern oysters (Crassostrea virginica). J Invertebr Pathol. 2008 Feb;97(2):150–8. doi: 10.1016/j.jip.2007.08.007. Epub 2007 Sep 1. PMID: 17931651.

8. Boudreau, S.A., & Worm, B. (2012). Ecological role of large benthic decapods in marine ecosystems: a review. Marine Ecology Progress Series, 469, 195–213.

9. Bruhn, J. B., Gram, L., & Belas, R. (2007). Production of antibacterial compounds and biofilm formation by Roseobacter species are influenced by culture conditions. Applied and environmental microbiology, 73(2), 442–450. 10.1128/AEM.02238-06

10. Brumbaugh, R. D., & Coen, L. D. (2009). Contemporary Approaches for Small-Scale Oyster Reef Restoration to Address Substrate Versus Recruitment Limitation: A Review and Comments Relevant for the Olympia Oyster, Ostrea lurida Carpenter 1864. J. of Shellfish Research, 28(1):147–161 (2009). 10.2983/035.028.0105

11. Burgess, C. M., Loeffler, A., & Fagan, P. K. (2017). High-throughput sequencing: A roadmap toward community ecology. Ecology and Evolution, 7(7), 2593–2601.

12. Caporaso JG, Lauber CL, Walters WA, Berg-Lyons D, Lozupone CA, Turnbaugh PJ, Fierer N, Knight R. 2011. Global patterns of 16S rRNA diversity at a depth of millions of sequences per sample. Proc Natl Acad Sci U S A 108:4516–4522. doi: 10.1073/pnas.1000080107.

13. Caporaso, J. G., Lauber, C. L., Walters, W. A., Berg-Lyons, D., Lozupone, C. A., Turnbaugh, P. J., Fierer, N., & Knight, R. (2012). Ultra-high-throughput microbial community analysis on the Illumina HiSeq and MiSeq platforms. The ISME Journal, 6(8), 1621–1624.

14. Chan, S.S.W., Wong, H.T., Thomas, M., Alleway, H.K., Hancock, B., Russell, B.D. (2022) Increased Biodiversity Associated With Abandoned Benthic Oyster Farms Highlight Ecosystem Benefits of Both Oyster Reefs and Traditional Aquaculture. Front. Mar. Sci. 9:862548. doi: 10.3389/fmars.2022.862548

15. Coen, L. D., & Luckenbach, M. W. (2000). Developing success criteria and goals for evaluating oyster reef restoration: Ecological function or resource exploitation? Ecological Engineering, 15(3-4), 323–343.

16. Coen, L. D., Brumbaugh, R. D., Bushek, D., Grizzle, R., Luckenbach, M. W., Posey, M. H., Powers, S. P., Tolley, S. G. (2007). Ecosystem services related to oyster restoration. Marine Ecology Progress Series, 341, 303–307.

17. Cyphert EL, Clare S, Dash A, et al. A Pilot Study of the Gut Microbiota in Spine Fusion Surgery Patients. HSS Journal®. 2023;0(0). doi:10.1177/15563316231201410

18. Dame, R.F., & Dame, R.F. (1996). Ecology of Marine Bivalves: An Ecosystem Approach (1st ed.). CRC Press. 10.1201/9781003040880

19. Dang H, Lovell CR. (2016) Microbial Surface Colonization and Biofilm Development in Marine Environments. Microbiol Mol Biol Rev; 80(1):91–138. doi: 10.1128/MMBR.00037-15. PMID: 26700108; PMCID: PMC4711185.

20. Diner, R. E., Zimmer-Faust, A., Cooksey, E., Allard, S., Kodera, S. M., Kunselman, E., Garodia, Y., Verhougstraete, M. P., Allen, A. E., Griffith, J., & Gilbert, J. A. (2023). Host and Water Microbiota Are Differentially Linked to Potential Human Pathogen Accumulation in Oysters. Applied and environmental microbiology, 89(7), e0031823. 10.1128/aem.00318-23

21. Dixon P. VEGAN, a package of R functions for community ecology. J Veg. Sci. 2003 Apr 9;14(6):927–30.

22. Estaki M, Jiang L, Bokulich N, McDonald D, González A, Kosciolek T, Martino C, Zhu Q, Birmingham A, Vázquez-Baeza Y, Dillon M, Bolyen E, Caporaso J, Knight R. QIIME 2 Enables Comprehensive End-to-End Analysis of Diverse Microbiome Data and Comparative Studies with Publicly Available Data. Curr Protoc Bioinformatics. 2020 Jun;70(1):e100.

23. Fernandes A, Reid J, Macklaim J, McMurrough T, Edgell D, Gloor G. Unifying the analysis of high-throughput sequencing datasets: characterizing RNA-seq, 16S rRNA gene sequencing and selective growth experiments by compositional data analysis. Microbiome. 2014; 2: 15.

24. Freeman, C. J., Gleason, D. F., & Kemp, D. W. (2016). Geomicrobiology of a Seagrass Sediment: A Combined Metagenomic and Geochemical Analysis of an Endangered Coastal Ecosystem. Frontiers in Microbiology, 7, 967.

25. Gatesoupe F (2008) Updating the importance of lactic acid bacteria in fish farming: natural occurrence and probiotic treatments. J Mol Microbiol Biotechnol 14:107–114

26. Gledhill, D. K., White, M. M., Salisbury, J., Thomas, H., Mlsna, I., Liebman, M., Mook, B., Grear, J., Candelmo, A. C., Chambers, R. C., Gobler, C. J., Hunt, C. W., King, A. L., Price, N. N., Signorini, S. R., Stancioff, E., Stymiest, C., Wahle, R. A., Waller, J. D., Rebuke, N.D., Wang, Z.A., Capson, T.L., Morrison, J.R., Cooley, S.R., Doney, S. C. (2015). Ocean and Coastal Acidification off New England and Nova Scotia. Oceanography, 28(2), 182–197. http://www.jstor.org/stable/24861880

27. Grabowski, J. H., Brumbaugh, R. D., Conrad, R. F., Keeler, A. G., Opaluch, J. J., Peterson, C. H., Piehler, M.F., Powers, S.P., Smyth, A.R. (2012). Economic valuation of ecosystem services provided by oyster reefs. BioScience, 62(10), 900–909.

28. Grabowski, J.H., Baillie, C.J., Baukus, A., Carlyle, R., Fodrie, F.J., Gittman, R.K., Hughes, A.R., Kimbro, D.L., Lee, Juhyung., Lenihan, H.S., Powers, S.P., Sullivan, K. (2022) Fish and invertebrate use of restored vs. natural oyster reefs in a shallow temperate latitude estuary. Ecosphere 13(5): e4035. 10.1002/ecs2.4035

29. Green TJ, Siboni N, King WL, Labbate M, Seymour JR, Raftos D. Simulated Marine Heat Wave Alters Abundance and Structure of Vibrio Populations Associated with the Pacific Oyster Resulting in a Mass Mortality Event. Microb Ecol. 2019 Apr;77(3):736–747. doi: 10.1007/s00248-018-1242-9. Epub 2018 Aug 10. PMID: 30097682.

30. Guedes, B.; Godinho, O.; Quinteira, S.; Lage, O.M. Antimicrobial Resistance Profile of PlanctomycetotaIsolated from Oyster Shell Biofilm: Ecological Relevance within the One Health Concept. Appl. Microbiol. 2024, 4, 16–26. 10.3390/applmicrobiol4010002

31. Harris, J. L., Deines, P., & Flint, H. J. (2016). The microbiome of the gastrointestinal tract of marine polychaetes: A review. Frontiers in Microbiology, 7, 2043.

32. Jiang Z., Du P., Liao Y., Liu Q., Chen Q., Shou L., et al. (2019). Oyster farming control on phytoplankton bloom promoted by thermal discharge from a power plant in a eutrophic, semi-enclosed bay. Water Res. 159, 1–9. doi: 10.1016/J.WATRES.2019.04.023

33. Kang CH, Gu T, So JS. Possible Probiotic Lactic Acid Bacteria Isolated from Oysters (Crassostrea gigas). Probiotics Antimicrob Proteins. 2018 Dec;10(4):728–739. doi: 10.1007/s12602-017-9315-5. PMID: 28875385.

34. Kellogg ML, Smyth AR, Luckenbach MW, Carmichael RH, Brown BL, Cornwell JC, Piehler MF, Owens MS, Dalrymple DJ, Higgins CB (2014). Use of oysters to mitigate eutrophication in coastal waters. Estuarine, Coastal and Shelf Science. 151:156–168.

35. King, G. M., Judd, C., Kuske, C. R., & Smith, C. (2012). Analysis of stomach and gut microbiomes of the Eastern Oyster (Crassostrea virginica) from coastal Louisiana, USA. PLOS ONE, 7(12), e51475.

36. Li PN, Herrmann J, Tolar BB, Poitevin F, Ramdasi R, Bargar JR, et al. Nutrient transport suggests an evolutionary basis for charged archaeal surface layer proteins. ISME J. 2018;12:2389–402.

37. Lokmer, A., & Wegner, K. M. (2015). Microbiome promotes tolerance to ocean acidification in oysters. Nature Climate Change, 5(5), 47–452.

38. Lu S, Liu X, Ma Z, Liu Q, Wu Z, Zeng X, Shi X and Gu Z (2016) Vertical Segregation and Phylogenetic Characterization of Ammonia-Oxidizing Bacteria and Archaea in the Sediment of a Freshwater Aquaculture Pond. Front. Microbiol. 6:1539. doi: 10.3389/fmicb.2015.01539

39. Luckenbach, M., Mann, R. L., & Wesson, J. A. (1999) Oyster Reef Habitat Restoration: a synopsis and synthesis of approaches; proceedings from the symposium, Williamsburg, Virginia, April 1995. Virginia Institute of Marine Science, William & Mary. 10.21220/V5NK51

40. Mallick H, Ma S, Franzosa EA, Vatanen T, Morgan XC, Huttenhower C. Experimental design and quantitative analysis of microbial community multiomics. Genome Biology. 2017 Nov 30;18(1):228.

41. Mallick H, Rahnavard A, McIver LJ, Ma S, Zhang Y, Nguyen LH, et al. Multivariable association discovery in population-scale meta-omics studies. PLoS Comput Biol. 2021; 17(11): e1009442.

42. Mannochio-Russo, H., Swift, S.O.I., Nakayama, K.K. et al. Microbiomes and metabolomes of dominant coral reef primary producers illustrate a potential role for immunolipids in marine symbioses. Commun Biol 6, 896 (2023). 10.1038/s42003-023-05230-1

43. Martens-Habbena W, Qin W, Horak REA, Urakawa H, Schauer AJ, Moffett JW, et al. The production of nitric oxide by marine ammonia-oxidizing archaea and inhibition of archaeal ammonia oxidation by a nitric oxide scavenger. Environ Microbiol. 2015;17:2261–74.

44. Morris, R. L., Bilkovic, D. M., Boswell, M. K., Bushek, D., Cebrian, J., Goff, J., Kibler, K. M., Peyre, M. K. L., McClenachan, G., Moody, J., Sacks, P., Shinn, J. P., Sparks, E. L., Temple, N. A., Walters, L. J., Webb, B. M., & Swearer, S. E. (2019). The application of oyster reefs in shoreline protection: Are we over-engineering for an ecosystem engineer? Journal of Applied Ecology, 56, 1703–1711.

45. Morris, R. L.; Bilkovic, Donna Marie; Boswell, M. K.; and et al, The application of oyster reefs in shoreline protection: Are we over-engineering for an ecosystem engineer? (2019). Journal of Applied Ecology, 56(7), 1703–1711. DOI: 10.1111/1365-2664.13390

46. Nearing JT, Douglas GM, Hayes MG, MacDonald J, Desai DK, Allward N, et al. Microbiome differential abundance methods produce different results across 38 datasets. Nat Commun. 2022; 13: 342.

47. Newell, R. I. E. (2004). Ecosystem influences of natural and cultivated populations of suspension-feeding bivalve molluscs: A review. Journal of Shellfish Research, 23(1), 51–61.

48. Newton IL, Woyke T, Auchtung TA, Dilly GF, Dutton RJ, Fisher MC, Fontanez KM, Lau E, Stewart FJ, Richardson PM, Barry KW, Saunders E, Detter JC, Wu D, Eisen JA, Cavanaugh CM. The Calyptogena magnifica chemoautotrophic symbiont genome. Science. 2007 Feb 16;315(5814):998–1000. doi: 10.1126/science.1138438. PMID: 17303757.

49. NOAA (2020). “Water cleaning capacity of oysters could mean extra income for chesapeake bay growers” Accessed on 02/14/2020 Retrieved from: https://coastalscience.noaa.gov/news/water-cleaning-capacity-of-oysters-could-mean-extra-income-for-chesapeake-bay-growers-video/

50. Okon, E.M.; Birikorang, H.N.; Munir, M.B.; Kari, Z.A.; Téllez-Isaías, G.; Khalifa, N.E.; Abdelnour, S.A.; Eissa, M.E.H.; Al-Farga, A.; Dighiesh, H.S.; et al. (2023) A Global Analysis of Climate Change and the Impacts on Oyster Diseases. Sustainability 15, 12775. 10.3390/su151712775

51. Qin, W., Zheng, Y., Zhao, F., Wang, Y., Urakawa, H., Martens-Habbena, W., Liu, H., Huang, X., Zhang, X., Nakagawa, T., Mende, D. R., Bollmann, A., Wang, B., Zhang, Y., Amin, S. A., Nielsen, J. L., Mori, K., Takahashi, R., Armbrust, E. V., Winkler, M.-K. H., DeLong, E. F., Li, M., Lee, P.-H., Zhou, J., Zhang, C., Zhang, T., Stahl, D. A., & Ingalls, A. E. (2020). Alternative strategies of nutrient acquisition and energy conservation map to the biogeography of marine ammonia-oxidizing archaea. The ISME Journal, 14(10), 2595–2609. 10.1038/s41396-020-0710-7

52. Remple, K.L., Silbiger, N.J., Quinlan, Z.A. et al. Coral reef biofilm bacterial diversity and successional trajectories are structured by reef benthic organisms and shift under chronic nutrient enrichment. npj Biofilms Microbiomes 7, 84 (2021). 10.1038/s41522-021-00252-1

53. Ridlon, A.D., Marks, A., Zabin, C.J. et al. Conservation of Marine Foundation Species: Learning from Native Oyster Restoration from California to British Columbia. Estuaries and Coasts 44, 1723–1743 (2021). 10.1007/s12237-021-00920-7

54. RStudio Team. RStudio: Integrated Development for R [Internet]. Boston, MA, USA: RStudio, PBC; 2021. Available from: http://www.rstudio.com

55. Sakowski EG, Wommack KE, Polson SW. Oyster Calcifying Fluid Harbors Persistent and Dynamic Autochthonous Bacterial Populations That May Aid in Shell Formation. Mar Ecol Prog Ser. 2020 Oct 29;653:57–75. doi: 10.3354/meps13487. PMID: 33424068; PMCID: PMC7789820.

56. Schulte DM, Burke RP, Lipcius RN. Unprecedented restoration of a native oyster metapopulation. Science. 2009 Aug 28;325(5944):1124–8. doi: 10.1126/science.1176516. Epub 2009 Jul 30. Erratum in: Science. 2010 Nov 5;330(6005):756. PMID: 19644073.

57. Scyphers, S. B., Powers, S. P., Heck, K. L. Jr., & Byron, D. (2011). Oyster Reefs as Natural Breakwaters Mitigate Shoreline Loss and Facilitate Fisheries. PLOS ONE, 6(8), e22396.

58. Smith, R.S., Cheng, S.L., Castorani, M.C.N. (2022) Meta-analysis of ecosystem services associated with oyster restoration. Conservation Biology 37(1): e13966. 10.1111/cobi.13966

59. Stevens JTE, Fulweiler RW, Roy Chowdhury P. 16S rRNA Amplicon Sequencing of Sediment Bacterial Communities in an Oyster Farm in Rhode Island. Microbiol Resour Announc. 2019 Oct 17;8(42):e01074–19. doi: 10.1128/MRA.01074-19. PMID: 31624174; PMCID: PMC6797539.

60. Stevick RJ, Sohn S, Modak TH, Nelson DR, Rowley DC, Tammi K, Smolowitz R, Markey Lundgren K, Post AF and Gómez-Chiarri M (2019) Bacterial Community Dynamics in an Oyster Hatchery in Response to Probiotic Treatment. Front. Microbiol. 10:1060. doi: 10.3389/fmicb.2019.01060

61. Thompson, L. R., Anderson, S. R., Den Uyl, P. A., Patin, N. V., Lim, S. J., Sanderson, G. & Goodwin, K. D. Tourmaline: A containerized workflow for rapid and iterable amplicon sequence analysis using QIIME 2 and Snakemake. GigaScience, Volume 11, 2022, giac066, 10.1093/gigascience/giac066.

62. Trabal Fernández, N., Mazón-Suástegui, J. M., Vázquez-Juárez, R., Ascencio-Valle, F., & Romero, J. (2014). Changes in the composition and diversity of the bacterial microbiota associated with oysters (Crassostrea corteziensis) during commercial production. FEMS Microbiology Ecology, 88(1), 69–83.

63. Trabal N, Mazón-Suástegui JM, Vázquez-Juárez R, Asencio-Valle F, Morales-Bojórquez E, Romero J. Molecular analysis of bacterial microbiota associated with oysters (Crassostrea gigas and Crassostrea corteziensis) in different growth phases at two cultivation sites. Microb Ecol. 2012 Aug;64(2):555–69. doi: 10.1007/s00248-012-0039-5. Epub 2012 Mar 27. PMID: 22450510.

64. V Perricone, M Mutalipassi, A Mele, M Buono, D Vicinanza, P Contestabile (2023), Nature-based and bioinspired solutions for coastal protection: an overview among key ecosystems and a promising pathway for new functional and sustainable designs, ICES Journal of Marine Science, 80(5), 1218–1239, 10.1093/icesjms/fsad080

65. Waldbusser GG, Hales B, Langdon CJ, Haley BA, Schrader P, Brunner EL, et al. (2015) Ocean Acidification Has Multiple Modes of Action on Bivalve Larvae. PLoS ONE 10(6): e0128376. 10.1371/journal.pone.0128376

66. Weiss S, Xu ZZ, Peddada S, Amir A, Bittinger K, Gonzalez A, Lozupone C, Zaneveld JR, Vázquez-Baeza Y, Birmingham A, Hyde ER, Knight R. Normalization and microbial differential abundance strategies depend upon data characteristics. Microbiome. 2017 Mar 3;5(1):27.

67. Welsh, R. M., Zaneveld, J. R., Rosales, S. M., Payet, J. P., Burkepile, D. E., & Thurber, R. V. (2015). Bacterial predation in a marine host-associated microbiome. The ISME Journal, 10(6), 1540–4 doi: 10.1038/ismej.2015.219.

68. Wilberg, M. J., Livings, M. E., Barkman, J. S., Morris, B. T., & Robinson, J. M. (2011). Overfishing, disease, habitat loss, and potential extirpation of oysters in for upper Chesapeake Bay, USA. Marine Ecology Progress Series, 436, 131–144.

69. Yu, L., Gan, J. (2021) Mitigation of Eutrophication and Hypoxia through Oyster Aquaculture: An Ecosystem Model Evaluation off the Pearl River Estuary. Environmental Science & Technology 2021 55 (8), 5506–5514 DOI: 10.1021/acs.est.0c06616

70. zu Ermgassen, P. S. E., Hancock, B., DeAngelis, B. M., Greene, J. K., Schuster, E., Spalding, M. D., & Brumbaugh, R. D. (2016). Setting objectives for oyster habitat restoration using ecosystem services: A manager’s guide. The Nature Conservancy, Arlington, VA.

71. Zu Ermgassen, P. S., Grabowski, J. H., Gair, J. R., & Powers, S. P. (2015). Quantifying Fish and Mobile Invertebrate Production from a threatened nursery habitat. Journal of Applied Ecology, doi: 10.1111/1365-2664.12576.

