## supplementarytables for "Combinatorial characterization of bacterial taxa-driven differences in the microbiome of oyster reefs"

**Supplementary Table 1.** Taxonomy of significant ASVs (p < 0.05; magnitude of fold change > |5|) in Biofilm relative to Oyster identified using ALDEx2.

| **ASV** | **Phylum** | **Class** | **Order** | **Family** | **Genus** | **Fold change** | **-log_10_ p-value** |
| --- | --- | --- | --- | --- | --- | --- | --- |
| daae43be6cf06991  f62a085ba8bff3b6 | Bacteroidetes | Bacteroidia | Bacteroidales | S24-7 |  | 9.69 | 1.74 |
| 4cbfff144d4e7a4e0  f4619ed505be070 | Proteobacteria | Gamma-proteobacteria | Enterobacteriales | Enterobacteriaceae |  | 9.12 | 1.61 |
| d114fb4c335125128  be28401522dd41a | Firmicutes | Bacilli | Lactobacillales | Streptococcaceae | Lactococcus | 8.78 | 1.65 |
| ca9c66d47347c03d  0343d6fe03ed8636 | Proteobacteria | Gamma-proteobacteria | Pasteurellales | Pasteurellaceae | Haemophilus | 8.78 | 1.66 |
| e708751e62639cc  0951c413f92ea7620 | Fusobacteria | Fusobacteriia | Fusobacteriales | Fusobacteriaceae | Fusobacterium | 8.63 | 1.62 |
| 45b76e49b6500d8  d245c2f27f6fbb143 | Firmicutes | Erysipelotrichi | Erysipelotrichales | Erysipelotrichaceae | Clostridium | 8.51 | 1.69 |
| 7b28c20e72c6c95b  3e604f0849245770 | Verrucomicrobia | Verrucomicrobiae | Verrucomicrobiales | Verrucomicrobiaceae | Akkermansia | 8.40 | 1.65 |
| f9c27600180b4963  044fe32e320254a4 | Bacteroidetes | Bacteroidia | Bacteroidales | S24-7 |  | 8.28 | 1.64 |
| 7c51b17f759d67b1  13474e5981aea92a | Bacteroidetes | Bacteroidia | Bacteroidales | S24-7 |  | 8.23 | 1.67 |
| 06f825b512d903b9  230e1a55d87359ee | Firmicutes | Bacilli | Lactobacillales | Streptococcaceae | Streptococcus | 7.86 | 1.61 |
| 6db820962a5bf3ba  7e0fe7f3f9dcc994 | Fusobacteria | Fusobacteriia | Fusobacteriales | Fusobacteriaceae | Fusobacterium | 7.86 | 1.53 |
| 9a13cbb8e1b1246  8bf22d6a05aafb0c0 | Bacteroidetes | Bacteroidia | Bacteroidales | Rikenellaceae |  | 7.82 | 1.52 |
| 0ad13b6d1cd98ad7  e8f6ee9909d33114 | Bacteroidetes | Bacteroidia | Bacteroidales | S24-7 |  | 7.34 | 1.49 |
| fae6d0c5cd2a5334  1a4ec984541b1541 | Fusobacteria | Fusobacteriia | Fusobacteriales | Fusobacteriaceae | Fusobacterium | 7.26 | 1.52 |
| 3562d3a0374b9f2e  d190c1a7aa7dedb7 | Firmicutes | Clostridia | Clostridiales | Lachnospiraceae | [Ruminococcus] | 7.26 | 1.54 |
| 65d43491988bfe55  7da4d86a5ba25dae | Firmicutes | Bacilli | Bacillales | Staphylococcaceae | Staphylococcus | 7.26 | 1.43 |
| 4f5efd25dacb5d63  9316e7291ff6ff8b | Proteobacteria | Beta-proteobacteria | Neisseriales | Neisseriaceae | Neisseria | 7.17 | 1.45 |
| 569653b1659271a  290facfcedd0de061 | Firmicutes | Bacilli | Lactobacillales | Enterococcaceae | Enterococcus | 7.17 | 1.48 |
| d46e2205f0c6ecf6  7b51f83d111c509c | Proteobacteria | Gamma-proteobacteria | Enterobacteriales | Enterobacteriaceae |  | 7.16 | 1.37 |
| 922d33a2963d3e87  a3d4df6b82371235 | Firmicutes | Bacilli | Bacillales | Staphylococcaceae | Staphylococcus | 7.15 | 1.45 |
| 4598e7db6dff8f01  9ff4f6c50dc05df4 | Firmicutes | Clostridia | Clostridiales | Lachnospiraceae | Dorea | 7.15 | 1.58 |
| 1328ddadac3f2d08  eb25e79a18f7e574 | Firmicutes | Clostridia | Clostridiales | Lachnospiraceae |  | 7.11 | 1.44 |
| 99deb3c5ecb022ec  05609ebd1112a557 | Bacteroidetes | Bacteroidia | Bacteroidales | Bacteroidaceae | Bacteroides | 7.05 | 1.50 |
| 0232b56c81bb04f  b94bdcfdf8b073ae1 | Bacteroidetes | Bacteroidia | Bacteroidales | Porphyromonadaceae | Porphyromonas | 6.85 | 1.31 |
| 811f358e4e89eecb  d75298ffce878cba | Firmicutes | Clostridia | Clostridiales | Lachnospiraceae |  | 6.79 | 1.33 |
| 08eba18fc9aa967e  5e29f20ee5041a60 | Firmicutes | Clostridia | Clostridiales | Lachnospiraceae |  | 6.64 | 1.43 |
| cdf14d2fed157f803  2715a22d3bf4573 | Bacteroidetes | Bacteroidia | Bacteroidales | Prevotellaceae | Prevotella | 6.46 | 1.41 |
| 8e74f6f63a5f9a304  aeb8284be71fd23 | Firmicutes | Clostridia | Clostridiales | Lachnospiraceae | Coprococcus | 6.46 | 1.42 |
| fb3376cd5dd7b9f4  c866e17253a3e5db | Firmicutes | Clostridia | Clostridiales | Lachnospiraceae | [Ruminococcus] | 6.44 | 1.34 |
| c75ae6008025d801  34bd14e9712f9d5c | Firmicutes | Clostridia | Clostridiales | Lachnospiraceae | Coprococcus | 6.40 | 1.40 |
| 11c6abaafa7ba53a  4d1a06ea9f7db98a | Firmicutes | Bacilli | Lactobacillales | Streptococcaceae | Streptococcus | 6.37 | 1.31 |
| 70dbac76545ca7f  0d29e40f47caf5706 | Bacteroidetes | Bacteroidia | Bacteroidales | [Paraprevotellaceae] | [Prevotella] | 6.35 | 1.33 |
| 54dcb911e3ab04e9  d5b30c3912100b41 | Firmicutes | Clostridia | Clostridiales |  |  | 6.28 | 1.40 |
| a5a7486c3e60ff3d  6cc20e9e6359e7ea | Firmicutes | Clostridia | Clostridiales | Lachnospiraceae |  | 6.23 | 1.35 |
| 8674dcf0cfe35939  037fa9a5b986866f | Fusobacteria | Fusobacteriia | Fusobacteriales | Fusobacteriaceae | Fusobacterium | 6.00 | 1.30 |
| 581a55014641e1cd  55cdc272c0365a28 | Firmicutes | Bacilli | Lactobacillales | Lactobacillaceae | Lactobacillus | 6.00 | 1.32 |
| bfa3561ca0d31bca  54be08e2d9f7937f | Firmicutes | Clostridia | Clostridiales | Lachnospiraceae |  | 5.99 | 1.38 |
| 28fa7b8781756830  ce1f51f3d8060df9 | Proteobacteria | Beta-proteobacteria | Burkholderiales | Burkholderiaceae | Burkholderia | 5.88 | 1.36 |
| 85c44c83eddc5d30  28261a1000b7d0e1 | Firmicutes | Bacilli | Gemellales | Gemellaceae |  | 5.87 | 1.34 |
| 958d78a02bef69a7  95806f97adb117ea | Firmicutes | Clostridia | Clostridiales | Ruminococcaceae | Oscillospira | 5.60 | 1.35 |
| f5e65944985d3999  1cba3fab949a0150 | Firmicutes | Clostridia | Clostridiales |  |  | 5.06 | 1.35 |
| a3e6c1e678f0a902  1dbf1b19a2766641 | Proteobacteria | Gamma-proteobacteria | Alteromonadales | OM60 |  | -5.37 | 1.61 |
| 8f2ac656dc37fe2bc  b4a8abe40ba1154 | Proteobacteria | Delta-proteobacteria | Desulfobacterales | Desulfobulbaceae |  | -5.42 | 1.33 |
| dce65a20a7cc132b  292aa8d8abd42d2d | Planctomycetes | Planctomycetia | Planctomycetales | Planctomycetaceae | Planctomyces | -6.25 | 2.25 |
| 74968ebe072e538f  b82e979d8986ad5b | Proteobacteria | Delta-proteobacteria | Desulfuromonadales | Desulfuromonadaceae |  | -6.68 | 1.56 |
| 06aa8371756490a  a26b512dac0839f29 | Bacteroidetes | [Saprospirae] | [Saprospirales] | Saprospiraceae |  | -7.44 | 2.72 |
| 094b90b37ab7cbb  5661727bdf30972b8 | Crenarchaeota | Thaumarchaeota | Cenarchaeales | Cenarchaeaceae | Nitrosopumilus | -8.65 | 3.04 |

**Supplementary Table 2.** Taxonomy of top 50 significant ASVs (p < 0.05; magnitude of fold change > |6|) in Biofilm relative to Sediment identified using ALDEx2.

| **ASV** | **Phylum** | **Class** | **Order** | **Family** | **Genus** | **Fold change** | **-log_10_ p-value** |
| --- | --- | --- | --- | --- | --- | --- | --- |
| daae43be6cf06991  f62a085ba8bff3b6 | Bacteroidetes | Bacteroidia | Bacteroidales | S24-7 |  | 9.91 | 1.79 |
| 4cbfff144d4e7a4e  0f4619ed505be070 | Proteobacteria | Gamma-proteobacteria | Enterobacteriales | Enterobacteriaceae |  | 9.37 | 1.88 |
| d114fb4c33512512  8be28401522dd41a | Firmicutes | Bacilli | Lactobacillales | Streptococcaceae | Lactococcus | 9.00 | 1.72 |
| ca9c66d47347c03d  0343d6fe03ed8636 | Proteobacteria | Gamma-proteobacteria | Pasteurellales | Pasteurellaceae | Haemophilus | 8.98 | 1.62 |
| e708751e62639cc0  951c413f92ea7620 | Fusobacteria | Fusobacteriia | Fusobacteriales | Fusobacteriaceae | Fusobacterium | 8.82 | 1.75 |
| 45b76e49b6500d8  d245c2f27f6fbb143 | Firmicutes | Erysipelotrichi | Erysipelotrichales | Erysipelotrichaceae | Clostridium | 8.54 | 1.65 |
| 6db820962a5bf3b  a7e0fe7f3f9dcc994 | Fusobacteria | Fusobacteriia | Fusobacteriales | Fusobacteriaceae | Fusobacterium | 8.47 | 1.67 |
| 7c51b17f759d67b1  13474e5981aea92a | Bacteroidetes | Bacteroidia | Bacteroidales | S24-7 |  | 8.46 | 1.62 |
| 7b28c20e72c6c95b  3e604f0849245770 | Verrucomicrobia | Verrucomicrobiae | Verrucomicrobiales | Verrucomicrobiaceae | Akkermansia | 8.26 | 1.58 |
| f9c27600180b4963  044fe32e320254a4 | Bacteroidetes | Bacteroidia | Bacteroidales | S24-7 |  | 8.25 | 1.58 |
| 9a13cbb8e1b1246  8bf22d6a05aafb0c0 | Bacteroidetes | Bacteroidia | Bacteroidales | Rickenellaceae |  | 8.17 | 1.62 |
| 06f825b512d903b9  230e1a55d87359ee | Firmicutes | Bacilli | Lactobacillales | Streptococcaceae | Streptococcus | 7.93 | 1.64 |
| 4f5efd25dacb5d63  9316e7291ff6ff8b | Proteobacteria | Beta-proteobacteria | Neisseriales | Neisseriaceae | Neisseria | 7.80 | 1.54 |
| 65d43491988bfe55  7da4d86a5ba25dae | Firmicutes | Bacilli | Bacillales | Staphylococcaceae | Staphylococcus | 7.66 | 1.50 |
| 569653b1659271a  290facfcedd0de061 | Firmicutes | Bacilli | Lactobacillales | Enterococcaceae | Enterococcus | 7.58 | 1.44 |
| 99deb3c5ecb022ec  05609ebd1112a557 | Bacteroidetes | Bacteroidia | Bacteroidales | Bacteroidaceae | Bacteroides | 7.44 | 1.52 |
| 922d33a2963d3e8  7a3d4df6b82371235 | Firmicutes | Bacilli | Bacillales | Staphylococcaceae | Staphylococcus | 7.42 | 1.63 |
| 1328ddadac3f2d08e  b25e79a18f7e574 | Firmicutes | Clostridia | Clostridiales | Lachnospiraceae |  | 7.37 | 1.50 |
| fae6d0c5cd2a53341  a4ec984541b1541 | Fusobacteria | Fusobacteriia | Fusobacteriales | Fusobacteriaceae | Fusobacterium | 7.31 | 1.46 |
| 0ad13b6d1cd98ad7e  8f6ee9909d33114 | Bacteroidetes | Bacteroidia | Bacteroidales | S24-7 |  | 7.31 | 1.58 |
| d46e2205f0c6ecf67  b51f83d111c509c | Proteobacteria | Gamma-proteobacteria | Enterobacteriales | Enterobacteriaceae |  | 7.28 | 1.44 |
| 3562d3a0374b9f2ed  190c1a7aa7dedb7 | Firmicutes | Clostridia | Clostridiales | Lachnospiraceae | [Ruminococcus] | 7.28 | 1.41 |
| 0232b56c81bb04fb  94bdcfdf8b073ae1 | Bacteroidetes | Bacteroidia | Bacteroidales | Porphyromonadaceae | Porphyromonas | 7.20 | 1.55 |
| 4598e7db6dff8f01  9ff4f6c50dc05df4 | Firmicutes | Clostridia | Clostridiales | Lachnospiraceae | Dorea | 7.15 | 1.35 |
| 08eba18fc9aa967e  5e29f20ee5041a60 | Firmicutes | Clostridia | Clostridiales | Lachnospiraceae |  | 7.04 | 1.50 |
| 811f358e4e89eecb  d75298ffce878cba | Firmicutes | Clostridia | Clostridiales | Lachnospiraceae |  | 6.99 | 1.49 |
| a5a7486c3e60ff3d6  cc20e9e6359e7ea | Firmicutes | Clostridia | Clostridiales | Lachnospiraceae |  | 6.85 | 1.48 |
| 11c6abaafa7ba53a4  d1a06ea9f7db98a | Firmicutes | Bacilli | Lactobacillales | Streptococcaceae | Streptococcus | 6.82 | 1.49 |
| fb3376cd5dd7b9f4c  866e17253a3e5db | Firmicutes | Clostridia | Clostridiales | Lachnospiraceae | [Ruminococcus] | 6.70 | 1.51 |
| c75ae6008025d801  34bd14e9712f9d5c | Firmicutes | Clostridia | Clostridiales | Lachnospiraceae | Coprococcus | 6.66 | 1.40 |
| f80166d86a7f15b69  b6ac97505299c3a | Bacteroidetes | Bacteroidia | Bacteroidales | S24-7 |  | 6.59 | 1.44 |
| 1fdb09b154f347c35  de3aad68774cb79 | Firmicutes | Clostridia | Clostridiales | Lachnospiraceae |  | 6.59 | 1.45 |
| 70dbac76545ca7f0  d29e40f47caf5706 | Bacteroidetes | Bacteroidia | Bacteroidales | [Paraprevotellaceae] | [Prevotella] | 6.49 | 1.52 |
| 8674dcf0cfe359390  37fa9a5b986866f | Fusobacteria | Fusobacteriia | Fusobacteriales | Fusobacteriaceae | Fusobacterium | 6.45 | 1.42 |
| e93b313352019b81  13e7fc66bf757917 | Firmicutes | Clostridia | Clostridiales | [Tissierellaceae] | Parvimonas | 6.44 | 1.40 |
| 8e74f6f63a5f9a304  aeb8284be71fd23 | Firmicutes | Clostridia | Clostridiales | Lachnospiraceae | Coprococcus | 6.44 | 1.34 |
| 28fa7b8781756830  ce1f51f3d8060df9 | Proteobacteria | Beta-proteobacteria | Burkholderiales | Burkholderiaceae | Burkholderia | 6.41 | 1.37 |
| 54dcb911e3ab04e9d  5b30c3912100b41 | Firmicutes | Clostridia | Clostridiales |  |  | 6.26 | 1.38 |
| bfa3561ca0d31bca  54be08e2d9f7937f | Firmicutes | Clostridia | Clostridiales | Lachnospiraceae |  | 6.23 | 1.37 |
| e147b59672d5b3fd  45f1892b61ca78e3 | Proteobacteria | Delta-proteobacteria | Desulfobacterales | Desulfobacteraceae | Desulfococcus | -6.32 | 1.49 |
| 8f2ac656dc37fe2bc  b4a8abe40ba1154 | Proteobacteria | Delta-proteobacteria | Desulfobacterales | Desulfobulbaceae |  | -6.33 | 2.64 |
| d961a76a1175c13d  8fedbf879b5741a7 | Proteobacteria | Gamma-proteobacteria | Chromatiales |  |  | -6.54 | 1.93 |
| 01628a01bc3bed5  6314f902c0b031542 | Proteobacteria | Gamma-proteobacteria | Thiotrichales | Piscirickettsiaceae |  | -6.57 | 1.88 |
| 3085613b4040ad7  af954b531bd49f1ce | Proteobacteria | Gamma-proteobacteria | Thiotrichales | Piscirickettsiaceae |  | -6.57 | 1.96 |
| 46a7475bfbd3a0d1  254bcf3bf02c594a | Proteobacteria | Gamma-proteobacteria |  |  |  | -6.57 | 1.93 |
| 094b90b37ab7cbb5  661727bdf30972b8 | Crenarchaeota | Thaumarchaeota | Cenarchaeales | Cenarchaeaceae | Nitrosopumilus | -6.62 | 1.38 |
| 1d4287101d4e69b6  bba45059df936480 | Proteobacteria | Delta-proteobacteria | Desulfobacterales | Desulfobulbaceae |  | -6.63 | 1.67 |
| e41ccff6b086bac68  198a11d750d1cc7 | Proteobacteria | Gamma-proteobacteria | Alteromonadales | OM60 |  | -6.68 | 2.09 |
| 74968ebe072e538f  b82e979d8986ad5b | Proteobacteria | Delta-proteobacteria | Desulfuromonadales | Desulfuromonadaceae |  | -7.89 | 1.91 |
| 4a02d2d2c0cab54a  95e3c0ebb9e3e9b9 | Proteobacteria | Gamma-proteobacteria | Thiotrichales | Piscirickettsiaceae |  | -8.37 | 2.95 |

**Supplementary Table 3.** Taxonomy significant ASVs (p < 0.05; magnitude of fold change > |1|) in Oyster relative to Sediment identified using ALDEx2.

| **ASV** | **Phylum** | **Class** | **Order** | **Family** | **Genus** | **Fold change** | **-log_10_ p-value** |
| --- | --- | --- | --- | --- | --- | --- | --- |
| 06aa8371756490aa  26b512dac0839f29 | Bacteroidetes | [Saprospirae] | [Saprospirales] | Saprospiraceae |  | 8.09 | 2.90 |
| 74968ebe072e538f  b82e979d8986ad5b | Proteobacteria | Delta-proteobacteria | Desulfuromonadales | Desulfuromonadaceae |  | -1.37 | 1.47 |
| b64810a9bcdbb39e  6ea6607148fa08ee | Proteobacteria | Gamma-proteobacteria | Thiotrichales | Piscirickettsiaceae |  | -4.45 | 1.56 |
| 55403e2e1f514243  54c46cdb9b8ca177 | Proteobacteria | Gamma-proteobacteria | [Marinicellales] | [Marinicellaceae] |  | -4.59 | 1.32 |
| 6d4a1f217388c16b1  e102634ddcc6bd8 | Bacteroidetes | [Saprospirae] | [Saprospirales] | Saprospiraceae |  | -4.68 | 1.69 |
| 0cdc00a78dbc33a33  42d98d02d3ee272 | Bacteroidetes | Flavobacteriia | Flavobacteriales | Flavobacteriaceae | Lutimonas | -4.71 | 1.70 |
| d3b10176a7be9b96  121e65037e7695be | Proteobacteria | Delta-proteobacteria | Syntrophobacterales |  |  | -4.79 | 1.71 |
| 41ba04645b4a16b1  2650f74d552af0a2 | Proteobacteria | Delta-proteobacteria | Desulfobacterales | Desulfobulbaceae |  | -5.47 | 1.65 |
| 240d24e7f48af450  00908f3af96da819 | Proteobacteria | Gamma-proteobacteria | Alteromonadales | OM60 |  | -5.48 | 1.77 |
| 01628a01bc3bed56  314f902c0b031542 | Proteobacteria | Gamma-proteobacteria | Thiotrichales | Piscirickettsiaceae |  | -5.74 | 1.70 |
| 06c35381364f5831  6a8fc367908d2b40 | Proteobacteria | Alpha-proteobacteria | Rhizobiales | Hyphomicrobiaceae |  | -6.05 | 2.01 |
| fb706aac74597b64  51e954ba734adca5 | Proteobacteria | Delta-proteobacteria | Desulfobacterales | Desulfobulbaceae |  | -6.11 | 1.38 |
| 65d8eb0e5bf45db9  a397fc4978b7eca7 | Proteobacteria | Alpha-proteobacteria | Rhizobiales | Hyphomicrobiaceae |  | -6.64 | 2.50 |
| e147b59672d5b3fd  45f1892b61ca78e3 | Proteobacteria | Delta-proteobacteria | Desulfuromonadales | Desulfobacteraceae | Desulfococcus | -6.65 | 1.77 |
| d0cebe17cac69e42  e48178849ab207c5 | Proteobacteria | Gamma-proteobacteria | Alteromonadales | OM60 |  | -7.00 | 2.56 |
| 4a02d2d2c0cab54a  95e3c0ebb9e3e9b9 | Proteobacteria | Gamma-proteobacteria | Thiotrichales | Piscirickettsiaceae |  | -8.30 | 3.06 |

**Supplementary Table 4.** Taxonomy significant ASVs (p < 0.05; magnitude of fold change > |1|) in Brickyard Park relative to Point Pinole identified using ALDEx2.

| **ASV** | **Phylum** | **Class** | **Order** | **Family** | **Genus** | **Fold change** | **-log_10_ p-value** |
| --- | --- | --- | --- | --- | --- | --- | --- |
| 43bf2e05902a9f38  eb400647f5b1e0a0 | Proteobacteria | Gamma-proteobacteria | Vibrionales | Vibrionaceae | Vibrio | 7.59 | 1.89 |
| e165ac749a8c091a  01064baaa327113b | Proteobacteria | Delta-proteobacteria | Desulfobacterales | Desulfobulbaceae |  | 7.57 | 1.77 |
| 84fdfa79b1ef84f84  f0d751c8f49bddd | Proteobacteria | Gamma-proteobacteria | Alteromonadales | OM60 |  | 7.18 | 1.86 |
| 094b90b37ab7cbb5  661727bdf30972b8 | Crenarchaeota | Thaumarchaeota | Crenarchaeales | Cenarchaeaceae | Nitrosopumilus | 6.27 | 1.31 |
| 8882141b9e9fdec8  55573b5f89c4934b | Proteobacteria | Gamma-proteobacteria | Alteromonadales | OM60 |  | 6.12 | 1.33 |
| 1f135f3b5c218fb49  5142276eaa99b23 | Bacteroidetes | Flavobacteriia | Flavobacteriales |  |  | 5.39 | 1.46 |
| c129bc28f5359caf  12978c7f01a229ee | Bacteroidetes | Cytophagia | Cytophagales | Flammeovirgaceae | Fulvivirga | 5.31 | 1.40 |
| 9860786036f54301  916be3ab54cbdd68 | Actinobacteria | Acidimicrobiia | Acidimicrobiales |  |  | 5.02 | 1.40 |

**Supplementary Table 4.** Taxonomy significant ASVs (p < 0.05; magnitude of fold change > |1|) in Brickyard Park relative to Heron’s Head identified using ALDEx2.

| **ASV** | **Phylum** | **Class** | **Order** | **Family** | **Genus** | **Fold change** | **-log_10_ p-value** |
| --- | --- | --- | --- | --- | --- | --- | --- |
| e165ac749a8c091a  01064baaa327113b | Proteobacteria | Delta-proteobacteria | Desulfobacterales | Desulfobulbaceae |  | 7.57 | 2.14 |
| 81b1d854fa98dcfc  7217f209ac736c09 | Planctomycetes | Planctomycetia | Pirellulales | Pirellulaceae | planctomycete | 6.69 | 1.72 |
| 2438c6bded651e17  749490fb3d360a95 | Proteobacteria | Alpha-proteobacteria | Rhodobacterales | Rhodobacteraceae | Roseovarius | 5.99 | 1.44 |
| 720f8a38dc4b7766  f4d076685c28addc | Proteobacteria | Delta-proteobacteria | Myxococcales | OM27 |  | 5.86 | 1.54 |
| 1f135f3b5c218fb4  95142276eaa99b23 | Bacteroidetes | Flavobacteriia | Flavobacteriales |  |  | 5.72 | 1.68 |

**Supplementary Table 5.** Taxonomy significant ASVs (p < 0.05; magnitude of fold change > |1|) in Brickyard Park relative to Dunphy Park identified using ALDEx2.

| **ASV** | **Phylum** | **Class** | **Order** | **Family** | **Genus** | **Fold change** | **-log_10_ p-value** |
| --- | --- | --- | --- | --- | --- | --- | --- |
| c129bc28f5359caf1  2978c7f01a229ee | Bacteroidetes | Cytophagia | Cytophagales | Flammeovirgaceae | Fulvivirga | 5.25 | 1.32 |
